# CellPilot: an agentic framework that pilots small language models through autonomous single-cell annotation

**DOI:** 10.64898/2026.07.06.736807

**Authors:** Siqi Jiang, Cong Qi, Yeqing Chen, Xun Song, Zhi Wei

**Affiliations:** Computer Science, New Jersey Institute of Technology, 154 Summit Street, Newark, 07102, New Jersey, USA

**Keywords:** Single-cell RNA sequencing, structured workflow orchestration, small language models, cell type annotation, autonomous analysis

## Abstract

Large language models can annotate cell types from marker gene lists, but they typically operate after preprocessing and clustering are complete, treating annotation as a terminal labeling step rather than controlling the analytical decisions that produce the evidence for cell identity. We present CellPilot, an agentic framework that guides a locally deployable small language model through the full single-cell analysis workflow, from raw count matrices to cluster-level annotation. CellPilot combines standard single-cell analysis tools with structured workflow control and observation-guided reasoning, allowing the model to plan analyses, execute tools, inspect intermediate results and revise decisions within a traceable session. On GTEx, structured workflow orchestration raised the same 8B model from 0.39 in a prompt-only setting to 0.89, closing most of the gap to GPT-4o (0.92) within the same framework; the framework gain was substantially larger for the smaller backbone across datasets (+0.35 versus +0.19). Across GTEx, Tabula Sapiens, and Mouse Cell Atlas, CellPilot achieves cluster-level annotation accuracies of 0.891, 0.750, and 0.773, outperforming representative reference-based, marker-based, and LLM-based methods. CellPilot confidence scores were associated with annotation correctness and supported post hoc filtering, while complete execution traces were retained for each analysis. These results suggest that structured workflow orchestration can be a critical determinant of performance in multi-step single-cell analysis, enabling locally deployable small language models to approach larger proprietary models while preserving transparency and practical usability.

## Introduction

Single-cell RNA sequencing (scRNA-seq) has transformed the study of cellular heterogeneity by enabling transcriptome-scale profiling of individual cells across tissues, developmental stages and disease contexts [1–3], motivating large-scale efforts to construct comprehensive cellular atlases of human tissues [4]. The analysis of these data is not a single classification step, but a sequence of dependent analytical decisions. Quality control determines which cells are retained; normalization and feature selection shape the representation of transcriptional variation; dimensionality reduction and clustering define the cell populations to be interpreted; and marker discovery provides the evidence used for cell type annotation. Because each step conditions the next, choices made early in the workflow can propagate to downstream biological interpretation [5, 6]. Cell type annotation is therefore a central output of scRNA-seq analysis, and its accuracy depends on the quality of the upstream workflow that produces the clusters and marker evidence.

A broad ecosystem of methods has been developed to improve the reproducibility and efficiency of this final annotation step [7–9]. Marker- and signature-based approaches, including SCINA, scCATCH and ScType, infer cell identity from curated marker genes or gene sets [10–12]. Reference-based methods such as SingleR, scmap and Cell BLAST map query profiles onto labeled atlases using correlation, nearest-neighbor or embedding-based similarity [13–15]. Supervised and deep-learning methods, including ACTINN, scPred, scDeepSort and transformer-based models, learn decision boundaries from annotated datasets [16–19]. Large-scale single-cell foundation models such as scGPT, scFoundation and CellPLM further extend this direction by pre-training on millions of cells across tissues and conditions, with complementary architectures exploring efficient long-context representation of single-cell profiles [20–22]. More recently, large language models (LLMs) have introduced a complementary direction by interpreting marker genes and tissue context through natural-language reasoning. GPTCelltype showed that LLMs can generate biologically plausible cell type labels from marker lists [23], and CASSIA extended this idea with structured reasoning chains for interpretable annotation [24]. These methods have advanced automated annotation, but most operate after preprocessing and clustering have already been completed. They annotate the output of an existing pipeline rather than controlling the analytical choices that produce that output. As a result, they cannot revise upstream decisions when marker evidence is ambiguous, clustering resolution is inappropriate or intermediate results indicate that the analysis should be adjusted.

This distinction is especially important for LLM-based analysis. Large proprietary models can be effective at interpreting biological language, but dependence on remote APIs raises concerns about cost, reproducibility and data governance in settings where genomic data must remain within institutional infrastructure [25, 26]. Open-source small language models, including recent releases of the Qwen series [27], provide a practical alternative for local deployment, but direct prompting is not sufficient for long, dependent analytical workflows. A language model asked to label marker genes in isolation has limited access to the evidence that determines whether those markers are reliable. Agentic frameworks offer a way to address this limitation by linking language-model reasoning to computational analyses, intermediate observations and the current analytical context, with mechanisms for tool invocation, chain-of-thought reasoning and verbal self-reflection [28–33]. Such frameworks have begun to support autonomous research workflows in chemistry [34] and clinical risk prediction [35], suggesting a similar potential for multi-step single-cell analysis. In single-cell analysis, the relevant question is therefore not only whether a language model can name a cell type from a gene list, but whether a structured workflow can guide a small local model through the analytical process that produces that gene list. We define this strategy as structured workflow orchestration, in which the current analytical context, established single-cell procedures and intermediate observations jointly guide decisions from data inspection to final annotation.

Here we present CellPilot, an agentic framework that guides a locally deployable small language model through the complete scRNA-seq analysis workflow, from raw count matrices to cluster-level annotation. CellPilot implements structured workflow orchestration by linking a Scanpy-based analytical toolkit with observation-guided reasoning [36]. During each analysis, the model selects analytical steps, evaluates intermediate results and revises subsequent decisions before producing final annotations. We focus on cell type annotation because it integrates the consequences of upstream quality control, feature selection, dimensionality reduction, clustering and marker discovery into a measurable biological task. Across GTEx, Tabula Sapiens and Mouse Cell Atlas [37–39], CellPilot achieved cluster-level annotation accuracies of 0.891, 0.750 and 0.773 under a unified hierarchical evaluation, exceeding representative reference-based, marker-based and LLM-based annotation methods under a common reference-cluster scoring protocol. These results show that the workflow produces accurate annotations across datasets with different tissue composition, species context and reference-label structure. In controlled backbone comparisons, the structured workflow substantially reduced dependence on model scale: within CellPilot, an open-source 8B model approached GPT-4o performance on GTEx, whereas the same model performed poorly in a prompt-only annotation setting. CellPilot also retained complete analytical records and produced confidence scores associated with annotation correctness, supporting prioritization for review. Structured workflow orchestration therefore provides a practical route for using locally deployable small language models in autonomous cell type annotation from raw single-cell data.

## Results

### CellPilot enables autonomous cell type annotation through structured workflow orchestration

CellPilot is an agentic framework for autonomous cell type annotation from raw single-cell RNA-sequencing data. Unlike LLM-based annotation approaches that assign labels after preprocessing and clustering have already been completed, CellPilot treats annotation as a workflow-level task whose accuracy depends on the analytical decisions that generate the evidence for cell identity. Starting from a raw count matrix, the agent coordinates quality control, normalization, feature selection, dimensionality reduction, clustering, marker discovery and final cluster-level cell type assignment within a single traceable session (Fig. 1). This design allows a small language model to reason over the same intermediate evidence that a human analyst would inspect when annotating single-cell data.

**Fig. 1.**
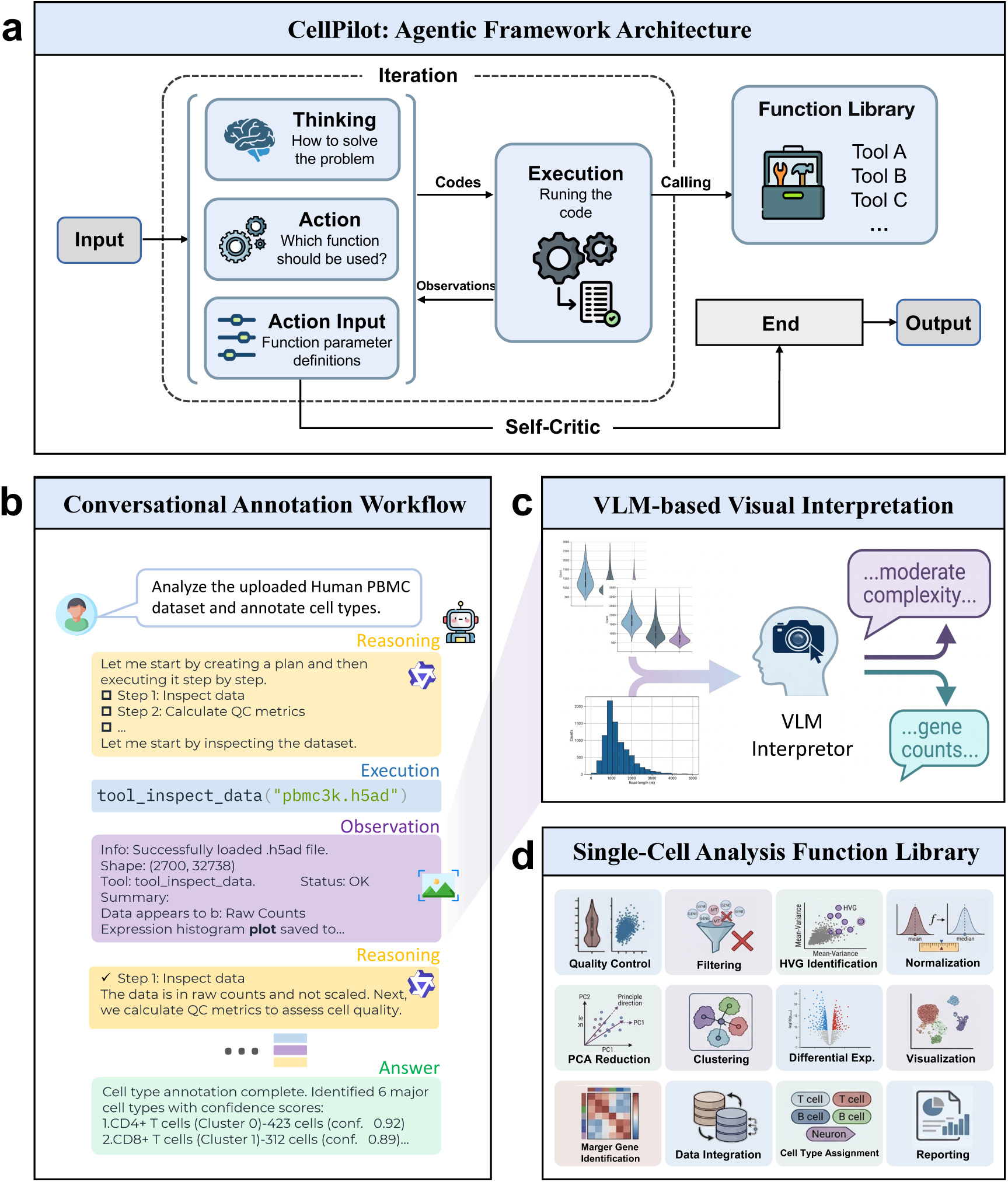
CellPilot: an agentic framework for autonomous single-cell annotation. a, Architecture of the CellPilot framework. The agent combines reasoning and tool execution through four components: *Thinking*(planning the strategy), *Action* (selecting a function), *Action Input* (specifying parameters), and *Execution* (running the tool). Results are returned as observations and evaluated by a *Self-Critic* module verifies that all required steps have been executed before the final output is produced. b, Conversational annotation workflow. Given a user request (eg., annotating a human PBMC dataset), the agent generates a stepwise plan, executes tools (for example `tool_inspect_data`), and reasons over observations in a dialogue-like loop to produce final cell-type annotations with confidence scores. c, VLM-based visual interpretation. Visualization outputs are interpreted by a vision–language model (VLM), which summarizes plot patterns and provides semantic descriptions that enrich the agent’s reasoning context. d, Single-cell analysis function library. The modular toolkit supports major stages of single-cell analysis, including quality control, filtering, highly variable gene identification, normalization, dimensionality reduction, clustering, differential expression, visualization, marker gene identification, data integration, cell-type assignment, and reporting.

Given an input expression matrix, CellPilot decomposes the annotation process into a sequence of state-dependent analytical decisions and executes them through an observation-driven reasoning loop (Fig. 1a,b). At each iteration, the agent selects an analytical function, specifies its parameters, executes the operation and incorporates the resulting observations into subsequent decisions. The function library covers the major stages required for annotation, including data inspection, quality control, filtering, normalization, highly variable gene selection, dimensionality reduction, clustering, differential expression, marker query, cell type assignment, visualization and reporting (Fig. 1d). A self-critic module monitors whether the required stages have been completed and returns control to the agent when additional steps are needed, reducing the risk of premature or incomplete annotation. CellPilot was designed to guide a locally deployable small language model through structured analytical evidence rather than depend on a large proprietary model for direct label generation. In this study, we used Qwen-8B as the default backbone, allowing the annotation workflow to be executed within local institutional infrastructure without dependence on remote model APIs (Methods). To incorporate visual evidence commonly used during manual annotation, CellPilot also includes a vision-language module that summarizes quality-control distributions, cluster layouts and marker-expression plots as textual observations for the agent context (Fig. 1c). These summaries provide auxiliary evidence for annotation decisions rather than serving as a separate annotation model.

We evaluated CellPilot across GTEx, Tabula Sapiens and Mouse Cell Atlas to test whether structured agent control enables a small language model to perform accurate cell type annotation from raw single-cell data. The following analyses examine the effect of the agent framework on backbone sensitivity, compare CellPilot with reference-based, marker-based and LLM-based annotation methods, and characterize the reliability, confidence and execution behavior of the resulting annotation workflow.

### Structured orchestration reduces dependence on backbone scale

A central question for autonomous cell type annotation is whether accurate performance requires a large proprietary language model, or whether a smaller locally deployable model can achieve comparable results when guided by a structured analytical workflow. We therefore compared Qwen3-8B and GPT-4o in two settings: within CellPilot, where the model reasons over workflow state, tool outputs and intermediate observations, and in GPTCelltype, where the same models perform annotation from marker-gene inputs without the CellPilot orchestration framework. Across GTEx, Tabula Sapiens and Mouse Cell Atlas, CellPilot showed only modest changes in mean hierarchical agreement when the backbone was switched from Qwen3-8B to GPT-4o, under both score-based and vote-based aggregation (Fig. 2a). The mean backbone gap, defined as |GPT-4o − Qwen3-8B|, was 0.056 for both CellPilot aggregation strategies, compared with 0.162 for GPTCelltype (Fig. 2b). Tissue-level paired comparisons showed the same pattern: CellPilot produced similar annotation scores across backbones, whereas GPTCelltype remained more sensitive to model scale, particularly in settings where Qwen3-8B had limited standalone annotation performance (Fig. 2c).

**Fig. 2.**
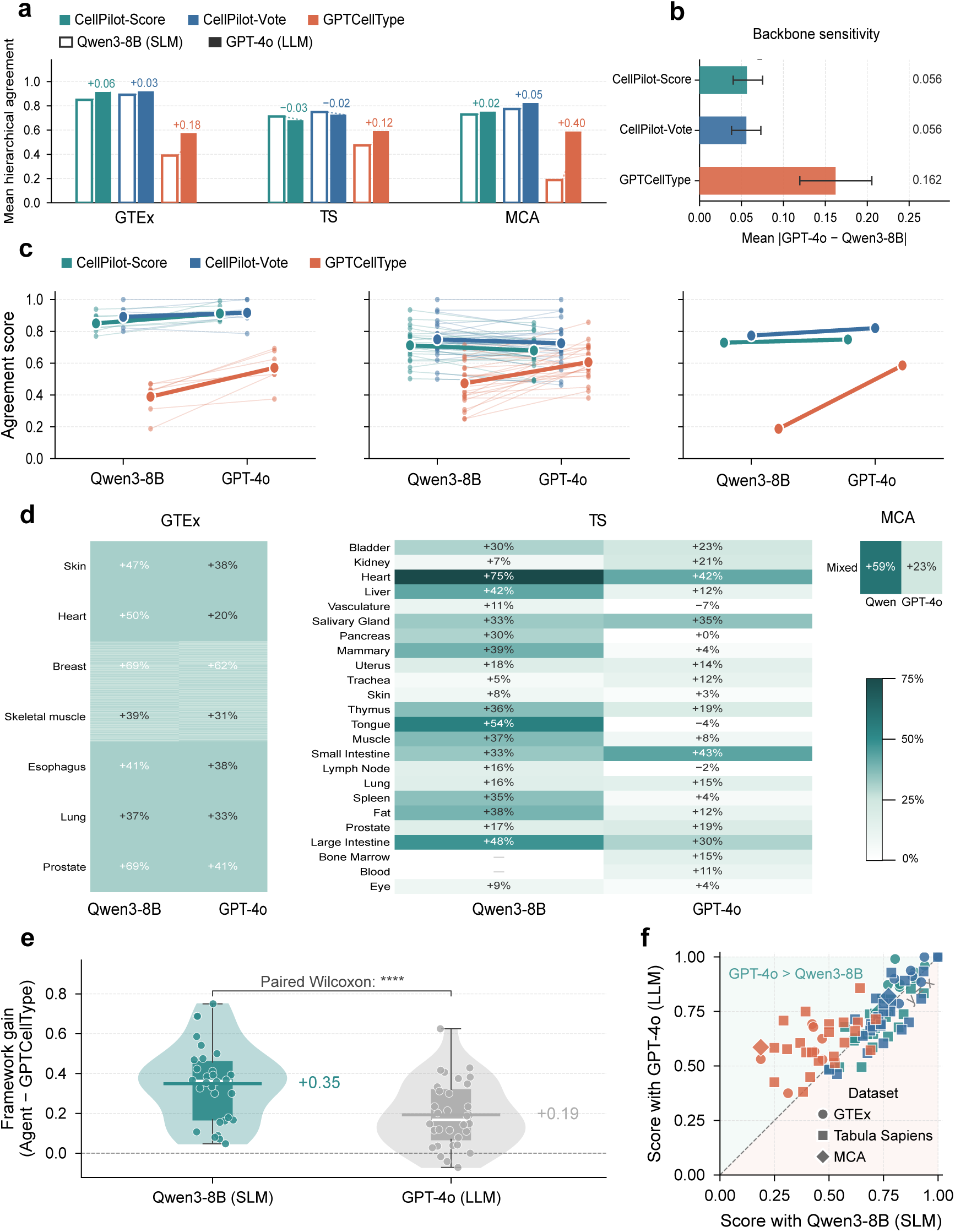
CellPilot’s agent framework decouples annotation performance from backbone capacity. **a**, Mean hierarchical agreement across three datasets (GTEx, Tabula Sapiens [TS], MCA) for CellPilot-Score, CellPilot-Vote and the framework-less baseline GPTCellType, each run with Qwen3-8B (hollow bars, SLM) and GPT-4o (filled bars, LLM). Numbers indicate the backbone gap Δ = GPT-4o − Qwen3-8B. **b**, Backbone sensitivity, defined as mean |Δ| across all (dataset, tissue) pairs. Error bars: 95% bootstrap CI (*n* = 2,000). **c**, Per-tissue slope of agreement score from Qwen3-8B to GPT-4o, by dataset. Thin lines: individual tissues; bold lines: per-method mean. **d**, Per-tissue structure gain Δ_struct_ = max(CellPilot) − GPTCellType under each backbone, in percentage points. **e**, Distribution of tissue-level structure gain by backbone, shown as violin + box + jittered points. Bars: mean. Paired Wilcoxon signed-rank test, *P <* 10*^−^*^4^ (∗∗∗∗). **f**, Per-tissue agreement with Qwen3-8B (*x*) versus GPT-4o (*y*). Shape: dataset; color: method. Points on *y* = *x* indicate identical SLM and LLM performance.

We next quantified how much performance was attributable to the framework itself by comparing CellPilot with GPTCelltype under the same backbone. The framework gain was positive across most tissues and datasets, indicating that structured orchestration improved annotation beyond direct marker-based prompting (Fig. 2d). This gain was larger for Qwen3-8B than for GPT-4o, with mean tissue-level improvements of 0.35 and 0.19, respectively (paired Wilcoxon signed-rank test, *P* < 10^-4^; Fig. 2e). Thus, the framework provided the greatest benefit when the underlying model had less standalone capacity, consistent with the idea that workflow structure can compensate for limitations in model scale.

Finally, per-tissue agreement between Qwen3-8B and GPT-4o showed that CellPilot narrowed the performance gap between small and large models while maintaining high annotation scores (Fig. 2f). These results indicate that CellPilot improves annotation not simply by replacing one language model with another, but by embedding the model within a constrained analytical process that supplies executable tools, intermediate biological evidence and explicit workflow state. In this setting, a locally deployable small model approached GPT-4o performance for cell type annotation.

### CellPilot produces accurate cell type annotations across benchmark datasets

We next evaluated whether CellPilot’s structured annotation workflow produces accurate cell type assignments across diverse single-cell datasets. Because cell type annotation depends on the preceding analytical steps that define clusters and marker genes, this comparison was designed to assess the complete annotation process rather than a single label-assignment prompt. We evaluated GTEx (7 tissues), Tabula Sapiens (24 tissues) and Mouse Cell Atlas (MCA) using the hierarchical scoring framework of Hou and Ji [23], in which fully correct, partially correct and incorrect predictions receive scores of 1, 0.5 and 0, respectively (Methods). CellPilot was compared with representative reference-based, marker-based and LLM-based annotation methods, including SingleR, ScType, GPTCelltype4 and CASSIA. These methods differ in their intended input assumptions: cluster-dependent baselines were evaluated using dataset-provided cluster information, whereas CellPilot started from raw count matrices and performed preprocessing, clustering, marker discovery and annotation within the same workflow. To make the comparison consistent, all predictions were scored against the same dataset-defined reference cell types.

CellPilot achieved the highest annotation scores across all three benchmark datasets (Fig. 3a). Results are reported using majority-vote aggregation (CellPilot-Vote). On GTEx, CellPilot reached a mean score of 0.891, compared with 0.584 for GPTCelltype4, 0.498 for CASSIA, 0.450 for ScType and 0.424 for SingleR (*P* < 0.001, Wilcoxon rank-sum test). On Tabula Sapiens, CellPilot achieved 0.750, exceeding GPTCelltype4 (0.589), CASSIA (0.569), ScType (0.489) and SingleR (0.446). On MCA, CellPilot achieved 0.773, compared with 0.508 for GPTCelltype4, 0.476 for CASSIA, 0.383 for SingleR and 0.273 for ScType. The improvement over reference- and marker-based methods was most pronounced in MCA, where cross-species annotation and uneven marker coverage pose challenges for methods that rely more directly on reference mapping or fixed marker dictionaries.

**Fig. 3.**
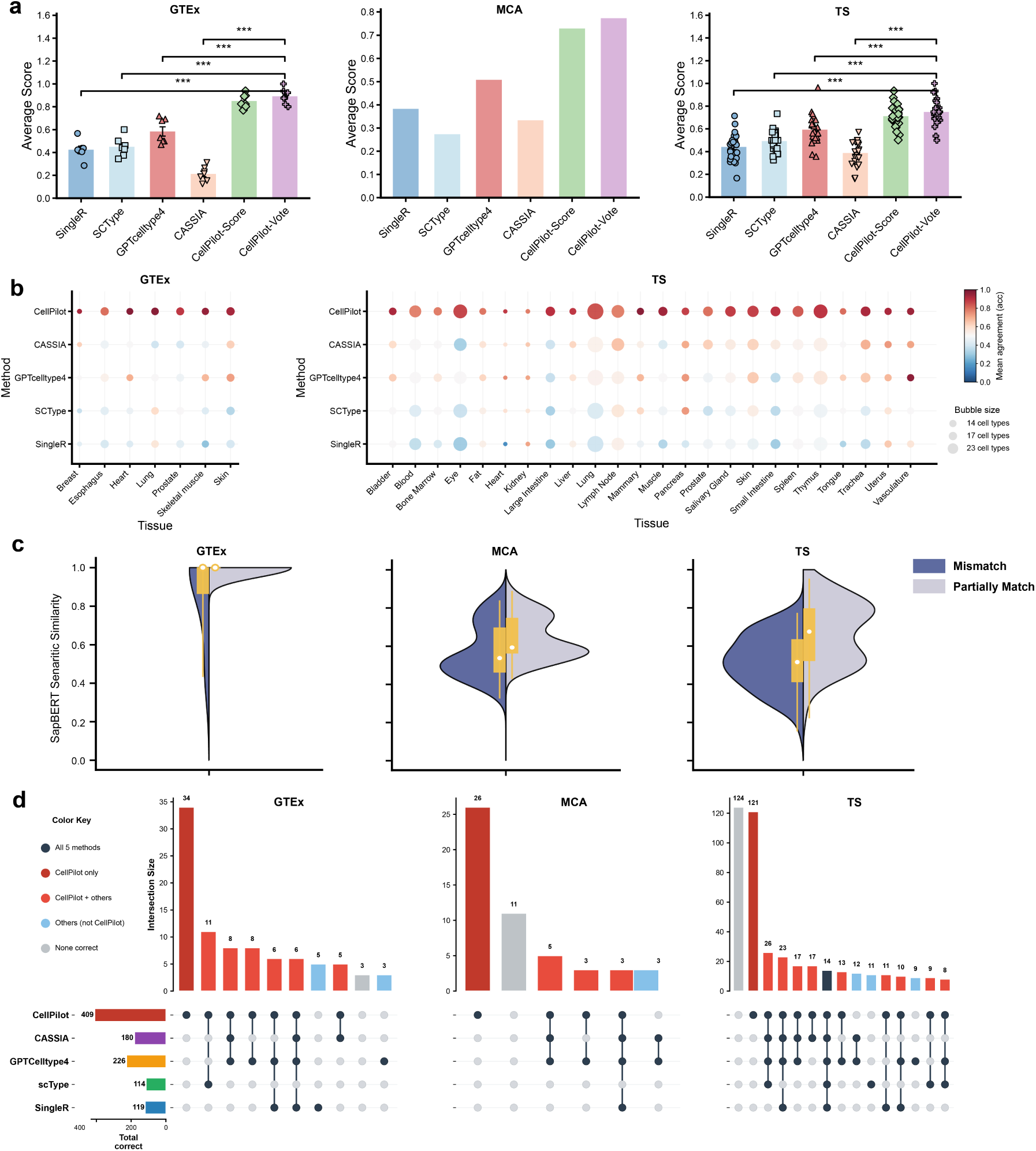
Benchmarking of CellPilot against baseline annotation methods. a, Average annotation scores for SingleR, ScType, GPTCelltype4, CASSIA, and CellPilot across GTEx, MCA, and Tabula Sapiens. Each point represents one tissue; for MCA only the dataset-level score is shown. Statistical significance was assessed by Wilcoxon rank-sum test (\*\*\**p <* 0.001). b, Tissue-level accuracy across GTEx and Tabula Sapiens. Bubble size indicates the number of cell types per tissue and color indicates mean agreement score. c, SapBERT cosine similarity between predicted and ground truth labels for partial matches (*S*(*c*) = 0.5) and mismatches (*S*(*c*) = 0) across all three datasets. Sample sizes (*n*) are indicated per group. d, UpSet plots of fully correct annotations (*S*(*c*) = 1, partial matches excluded) across GTEx, MCA, and Tabula Sapiens. Colors indicate annotation overlap patterns: CellPilot only (dark red), CellPilot with others (light red), all five methods (dark gray), others only (blue), and none correct (light gray). Horizontal bars show total correct annotations per method.

CellPilot’s performance was also consistent across individual tissues rather than being driven by a small number of favorable cases (Fig. 3b). In GTEx, CellPilot ranked highest across all seven tissues, maintaining high agreement across tissues with different cellular compositions. In Tabula Sapiens, CellPilot performed strongly across both immune-rich tissues, such as Blood and Bone Marrow, and more heterogeneous epithelial tissues, such as Lung and Large Intestine. Competing methods showed larger tissue-to-tissue variation, particularly in tissues with broader cellular diversity or closely related cell states. These results indicate that CellPilot’s annotation gains are not restricted to a single atlas or tissue context, but are preserved across human and mouse datasets with different annotation granularity, tissue composition and reference-label structure.

### CellPilot annotations remain biologically coherent beyond exact agreement

Hierarchical annotation scores provide a standardized measure of agreement, but they do not fully capture the biological distance between an incorrect prediction and the reference label. This distinction is important in cell type annotation, where labels often differ in granularity even when they refer to closely related identities within the Cell Ontology hierarchy. We therefore examined whether non-exact CellPilot annotations were biologically close to the reference labels by computing SapBERT cosine similarity for predictions scored as partial matches or mismatches (Fig. 3c).

Across GTEx, MCA and Tabula Sapiens, CellPilot predictions that did not receive full credit showed higher semantic similarity to the reference labels than the corresponding predictions from competing methods. This pattern was observed for both partial matches and complete mismatches, indicating that CellPilot errors were more often near misses within related cell identities rather than unrelated label assignments. For example, a prediction of “CD4^+^ T cells” for a reference label of “T helper cells” is penalized under exact ontology agreement, but still reflects a closely related immune-cell identity. Similar trends were observed when each baseline method was examined separately (Supplementary Fig. S1–S3), suggesting that CellPilot’s annotation behavior remains biologically coherent even when the predicted label does not exactly match the reference term.

We next asked whether CellPilot primarily recovered the same annotations as existing methods or contributed correct assignments not captured by other approaches. UpSet analysis of fully correct annotations showed that CellPilot accounted for the largest number of uniquely correct labels across the three datasets (Fig. 3d). CellPilot produced 409 annotations that were not correctly assigned by any other method, compared with 226 for GPTCelltype4, 180 for CASSIA, 119 for SingleR and 114 for ScType. Thus, CellPilot’s performance was not solely driven by agreement with existing annotation strategies or by partial matches within the scoring scheme. Instead, the method resolved a substantial set of cell type labels that were missed by reference-based, marker-based and prompt-based LLM approaches, supporting the value of coordinating annotation with upstream single-cell analysis rather than treating label assignment as an isolated step.

### Cell-level annotation performance varies with annotation granularity and cluster size

After establishing the overall cluster-level accuracy of CellPilot, we examined how annotation performance varied across individual cells, label granularity and cluster size, which together determine the strength and specificity of the evidence available for cell type assignment. Across 694,701 annotated cells in GTEx and Tabula Sapiens, 79.2% of cells received fully matched annotations, 14.5% received partially matched annotations and 6.3% were mismatched (Fig. 4a). These results indicate that most cells were assigned labels that agreed with the reference annotation under the Cell Ontology hierarchy.

**Fig. 4.**
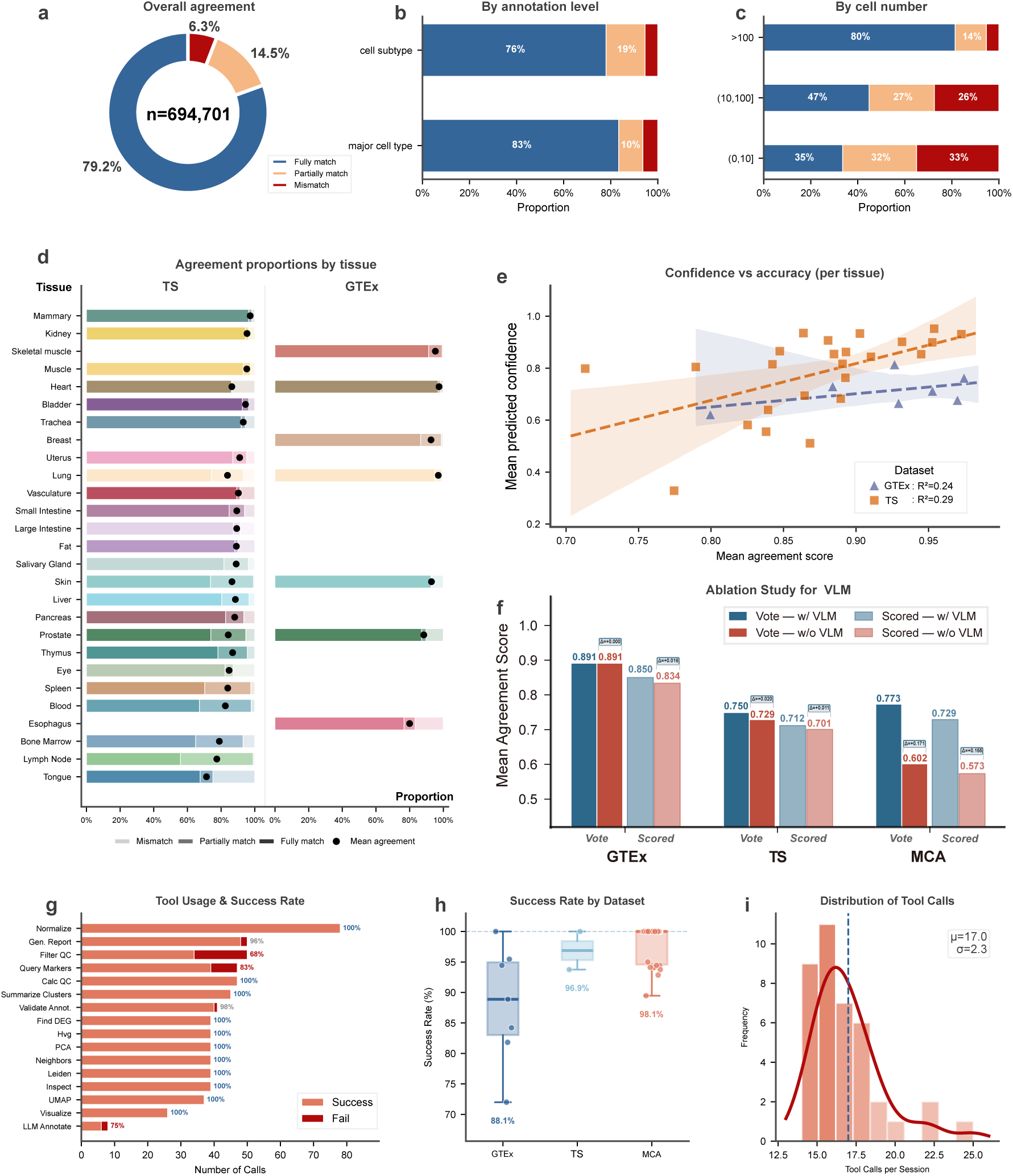
Cell-level annotation performance and operational reliability of CellPilot. a, Overall agreement proportions across all annotated cells, showing the fraction of fully matched, partially matched, and mismatched predictions. b, Agreement proportions stratified by annotation granularity, comparing performance on major cell types versus cell subtypes. c, Agreement proportions stratified by cluster size across three categories: small (0, 10], medium (10, 100], and large (*>*100) cells per cluster. d, Agreement proportions by tissue for Tabula Sapiens and GTEx. Each bar shows the proportion of fully matched, partially matched, and mismatched annotations, with black dots indicating mean agreement score. e, Correlation between mean predicted confidence *C*(*c*) and mean agreement score Acc_tissue_ per tissue for GTEx (*R*^2^ = 0.24) and Tabula Sapiens (*R*^2^ = 0.29), with 95% confidence intervals shown as shaded regions. f, Distribution of predicted confidence *C*(*c*) across fully matched, partially matched, and mismatched predictions. g, Per-tool call frequency and success rate across all sessions, with colors distinguishing successful (orange) and failed (red) executions. h, Overall workflow success rate *R* by dataset. i, Distribution of tool calls per session (*µ* = 17.0, *σ* = 2.3).

Annotation accuracy was influenced by the biological resolution of the reference labels. Major cell types were annotated with a full-match rate of 83%, compared with 76% for cell subtypes (Fig. 4b), consistent with the greater marker overlap and narrower transcriptional boundaries that distinguish closely related subpopulations. To assess whether errors were concentrated within particular lineages, we further mapped labels to broad cell type categories and examined the ten most frequent categories in human and mouse datasets (Supplementary Note 7, Supplementary Figs. 4–5). Structurally well-defined lineages, including fibroblasts and endothelial cells, showed consistently high agreement, whereas more transcriptionally overlapping populations, such as myeloid and macrophage/monocyte categories, showed higher rates of within-lineage confusion. Thus, most broad-category errors reflected ambiguity among related cell identities rather than arbitrary reassignment across unrelated lineages.

Cluster size provided an additional source of variation in annotation quality. Large clusters containing more than 100 cells achieved an 80% full-match rate, whereas medium clusters containing 10–100 cells and small clusters containing 10 or fewer cells achieved full-match rates of 47% and 35%, respectively (Fig. 4c). This pattern is expected because small clusters provide weaker differential-expression evidence and less stable marker-gene rankings for annotation. At the tissue level, agreement proportions also varied across organ systems, with immune-rich tissues such as Bone Marrow and Lymph Node showing high full-match rates, whereas tissues with more heterogeneous or glandular compositions showed higher proportions of partial matches (Fig. 4d). Consistent with this interpretation, tissue-level cell type diversity was only weakly associated with mean agreement but showed a stronger negative association with predicted confidence (Supplementary Fig. 6). These analyses define the operating characteristics of CellPilot: annotation is most reliable when cluster-level marker evidence is strong, whereas small or compositionally complex cell populations should be prioritized for review.

### Confidence scores and visual evidence support annotation assessment

Because autonomous annotation should indicate which predictions warrant additional inspection, we next assessed whether CellPilot confidence scores reflected annotation reliability. CellPilot assigns each cluster annotation a confidence score based on the relative support of marker genes for the predicted identity (Methods). At the tissue level, mean predicted confidence was positively associated with mean agreement score in both GTEx (*R*^2^ = 0.24) and Tabula Sapiens (*R*^2^ = 0.29; Fig. 4e). Although these correlations do not imply perfect calibration, they indicate that confidence scores capture useful variation in annotation quality across tissues. Calibration analysis further showed a tendency toward underconfidence at intermediate score ranges, with observed full-match rates often exceeding predicted confidence (Supplementary Fig. 7a). Confidence distributions differed across datasets, particularly in MCA, where broader and more multimodal scores reflected greater variation in cluster size and cell type composition (Supplementary Fig. 7b).

We further evaluated the practical use of confidence scores by examining the trade-off between annotation accuracy and retained coverage across confidence thresholds. Increasing the confidence threshold enriched the retained annotations for fully matched predictions while reducing the number of clusters retained for automatic acceptance (Supplementary Fig. 7c). This behavior supports the use of confidence as a triage signal rather than as an absolute probability of correctness: high-confidence annotations can be accepted with greater assurance, whereas low-confidence or small-cluster annotations can be directed to manual review or additional validation.

Finally, we assessed whether visual observations contributed additional information during annotation. CellPilot uses a vision-language module to summarize diagnostic plots, including quality-control distributions, cluster layouts and marker-expression visualizations, as textual observations for the agent context. Ablation analysis showed that the effect of this module was dataset dependent (Fig. 4f). In GTEx, vote-based accuracy was unchanged, while score-based agreement increased from 0.834 to 0.850. In Tabula Sapiens, vote-based agreement increased from 0.729 to 0.750 and score-based agreement from 0.701 to 0.720. The largest improvement occurred in MCA, where vote-based agreement increased from 0.602 to 0.773 and score-based agreement from 0.573 to 0.729. Tissue-level analyses showed that visual observations provided gains in several heterogeneous tissues but had little effect, and occasionally modest negative effects, in tissues with clearer transcriptional separation (Supplementary Fig. 8). These results indicate that confidence scores and visual observations are most useful as supporting signals for annotation review and refinement, particularly when marker evidence or cluster structure is ambiguous.

### CellPilot executes multi-step workflows with traceable and recoverable behavior

A practical annotation agent must not only produce accurate labels, but also complete the analytical steps that lead to those labels without requiring manual intervention. We therefore examined whether CellPilot could execute the full annotation workflow reliably across benchmark sessions, from raw data inspection to final cell type assignment and reporting. Workflow completion rates were high across datasets, reaching 98.1% in Tabula Sapiens and 96.9% in MCA, with a lower but still robust completion rate of 88.1% in GTEx (Fig. 4h). The lower GTEx rate reflected greater heterogeneity in input formatting and data-quality conventions across tissue contributions, rather than a single recurrent failure mode. Across all completed sessions, CellPilot required a mean of 17.0 ± 2.3 tool calls, indicating that the agent followed a bounded sequence of analytical steps rather than repeatedly invoking tools without convergence (Fig. 4i).

Step-level analysis showed that most core single-cell operations were executed with high reliability (Fig. 4g). Deterministic analytical steps, including normalization, highly variable gene selection, dimensionality reduction, neighborhood graph construction, clustering, visualization and report generation, succeeded in nearly all sessions. Steps that depend more strongly on dataset-specific properties showed lower success rates. Quality-control filtering was less stable when datasets had unusual metric distributions that did not match standard filtering heuristics, and marker-query failures were concentrated in rare or poorly characterized populations with limited marker database coverage. The LLM annotation step was invoked only for clusters that remained unresolved after marker-based matching; failures in this step therefore reflected ambiguous marker evidence rather than failure of the primary annotation path.

Importantly, individual tool failures did not necessarily terminate the annotation workflow. When a tool call failed, CellPilot received the error as an observation and used it to revise the subsequent analysis, either by adjusting parameters, returning to an earlier step or selecting an alternative operation. Clusters that remained unresolved after marker-based and LLM-assisted annotation were explicitly flagged for review rather than assigned a low-confidence label without indication. This behavior is central to the design of CellPilot: annotation decisions are accompanied by intermediate observations, confidence information and recoverable execution traces. Together, these results show that CellPilot functions as a traceable annotation workflow rather than a single-pass labeling prompt, allowing both successful assignments and unresolved cases to be inspected after execution.

### Case study: annotating a keratinocyte transformation trajectory from normal skin to actinic keratosis and cutaneous squamous cell carcinoma

Single-cell RNA sequencing has been widely applied to characterize the cellular ecosystems of cutaneous and head-and-neck squamous tumours, revealing extensive intra-tumoural cell-state heterogeneity that challenges manual annotation [40, 41]. To evaluate whether autonomous cell-type annotation can support biologically meaningful interpretation in such settings, we applied CellPilot to a publicly available single-cell RNA sequencing dataset spanning normal skin (NS), actinic keratosis (AK), and cutaneous squamous cell carcinoma (SCC) [42]. The dataset comprises nine samples with three biological replicates per condition and presents a challenging setting with heterogeneous cellular composition, disease-associated transcriptional remodeling, and no ground-truth annotations. Each sample was processed independently using our Qwen-8B-powered agent without manual curation, followed by integration using PCA and Harmony [43] for batch correction in a shared UMAP space. The integrated embedding revealed coherent epithelial, stromal, vascular, and immune compartments that were consistently recovered across independently processed samples (Fig. 5a).

**Fig. 5.**
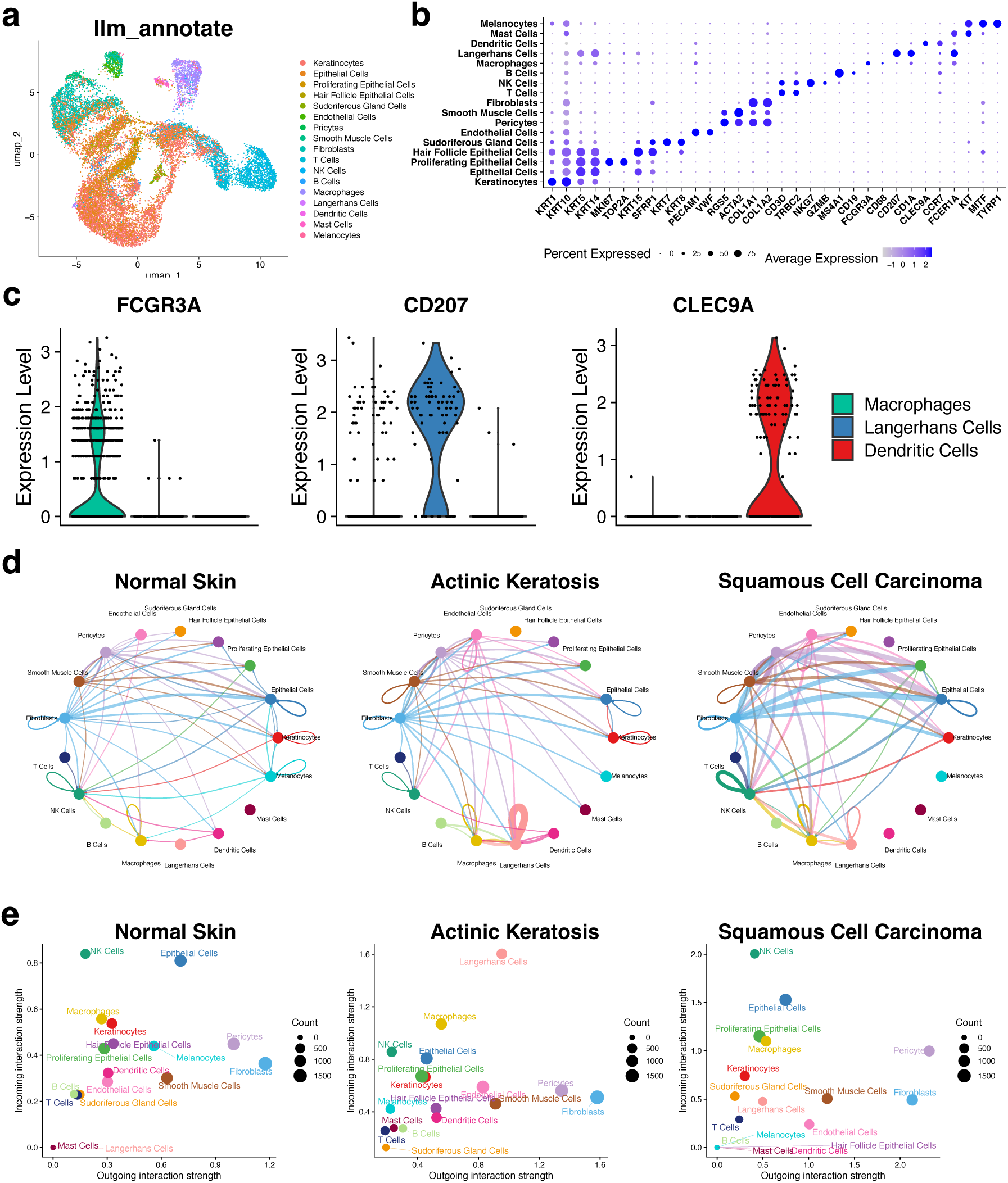
Integrated cell-type annotation and downstream cell–cell communication analysis across normal skin, actinic keratosis, and squamous cell carcinoma. a, UMAP of all cells integrated across nine samples, colored by cell types annotated by CellPilot. b, Dot plot showing the expression of selected canonical marker genes for each annotated cell type. Two representative marker genes are displayed per cell type. Dot size indicates the percentage of cells expressing the gene and color intensity represents average expression level. c, Violin plots showing the expression of representative marker genes for macrophages, Langerhans cells, and dendritic cells, highlighting distinct expression patterns among these closely related immune populations. d, Circle plots depicting cell–cell communication networks inferred by CellChat for each condition. Only the top 20% of interactions ranked by interaction strength are shown. Node colors represent cell types, edge thickness corresponds to interaction strength, and arrow direction indicates the direction of signaling. e, Scatter plots of signaling roles inferred by CellChat showing the relative outgoing and incoming interaction strengths of each cell type per condition.

Annotation validity was first assessed through canonical marker expression, which showed strong cell-type specificity with minimal off-target signal (Fig. 5b). Beyond major cell populations, the agent autonomously identified a rare Langerhans cell population that was not explicitly included in the provided annotation prior. Importantly, the agent was initialized only with common skin cell markers and was not supplied with Langerhans-specific marker genes, curated references, or predefined label vocabularies containing this population. The inferred identity was subsequently validated through examination of lineage-specific markers, including CD207, which showed mutually exclusive expression relative to neighboring immune populations such as macrophages and dendritic cells (Fig. 5c).

This result highlights a key limitation of conventional annotation workflows. Most existing approaches rely on predefined references, curated marker lists, or closed label spaces, and therefore primarily recover cell identities already represented in prior knowledge. In contrast, the agent was able to propose a biologically valid but non-obvious population outside the explicitly provided annotation space. Notably, Langerhans cells were embedded within a closely related immune transcriptional neighborhood, making their identification nontrivial even for experienced analysts without domain-specific prior familiarity. Rather than reflecting only increased clustering granularity, this finding demonstrates that autonomous agentic reasoning can expand the candidate biological hypothesis space beyond the analyst’s initial assumptions.

We next assessed whether these annotations supported downstream systems-level analysis. Reconstruction of cell–cell communication networks using CellChat [44] across NS, AK, and SCC revealed disease-associated interaction remodeling consistent with progressive epithelial transformation (Fig. 5d). Together, these results suggest that agent-derived annotations are biologically coherent, reproducible across independently processed samples, and sufficiently robust to support downstream analyses without manual curation.

## Discussion

CellPilot was developed for autonomous cell type annotation from raw single-cell RNA-sequencing data. Its central premise is that annotation should not be treated as a terminal label-assignment step, because the evidence used to assign cell identity is shaped by upstream choices in quality control, normalization, feature selection, clustering and marker discovery. CellPilot links these decisions within a state-machine-controlled agentic workflow, allowing a locally deployable small language model to inspect intermediate results, execute single-cell analysis tools and revise subsequent decisions before producing cluster-level annotations. This design differs from prompt-based annotation methods that operate on marker lists after preprocessing and clustering have already been completed.

The main methodological finding is that workflow orchestration can substantially reduce dependence on backbone model scale in autonomous cell type annotation. Under direct prompt-based annotation, Qwen3-8B remained markedly behind GPT-4o, whereas within CellPilot the performance gap between the two models was much smaller across the three benchmark datasets under both score-and vote-based aggregation. The framework gain was also larger for Qwen3-8B than for GPT-4o, indicating that orchestration is most beneficial when the underlying model has limited standalone annotation capacity. Accurate annotation, in this setting, does not arise simply from using a larger language model. It is supported by constraining model reasoning with workflow context, executable tools and biological observations generated during the analysis.

CellPilot produced accurate annotations across GTEx, Tabula Sapiens and Mouse Cell Atlas, outperforming representative reference-based, marker-based and LLM-based annotation methods under a unified hierarchical evaluation. These methods represent distinct strategies for automated annotation: mapping query profiles to reference atlases, matching marker signatures or using language models to interpret marker genes. CellPilot addresses the same task from a different angle, by coordinating the upstream analysis that produces the clusters and marker evidence used for label assignment. Its advantage was evident not only in aggregate agreement but also in the behavior of non-exact predictions. Semantic analysis showed that CellPilot annotations that did not exactly match the reference labels were often close to the correct identity within the Cell Ontology hierarchy, and UpSet analysis showed that CellPilot contributed a large number of uniquely correct annotations. Thus, the framework improved annotation quality beyond increasing the number of exact matches, yielding errors that were more biologically coherent and resolving labels missed by existing strategies.

The practical value of CellPilot depends on more than annotation accuracy. Autonomous annotation should also expose uncertainty, recover from failures and leave an analytical record that can be inspected after execution. CellPilot addresses these requirements through confidence scores, structured observations and recoverable tool execution. The confidence scores were associated with annotation correctness and supported prioritization of predictions for review, although they should not be interpreted as perfectly calibrated probabilities. Transparent reporting of uncertainty and execution traces is consistent with emerging reporting standards for studies that use large language models in biomedical settings [45]. The vision-language module provided additional visual evidence in a context-dependent manner, with the largest gains in MCA and smaller or negligible effects in several human tissues. Confidence estimation and visual interpretation therefore serve as supporting mechanisms for annotation assessment, rather than independent sources of method novelty.

Several limitations remain. CellPilot performed less well on very small clusters, where differential-expression evidence is unstable and marker-gene rankings are less reliable. This limitation is not specific to language models, but reflects a broader challenge in annotating rare or sparsely sampled populations from weak cluster-level evidence. Pan-cancer analyses have further shown that intra-tumoural cell-state heterogeneity is a pervasive feature across tumour types [46], motivating annotation methods that remain robust under such variation. Performance was also lower on MCA than on the human datasets, consistent with the greater heterogeneity of the mouse atlas and less complete marker coverage for some mouse cell types. Incorporating species-specific marker resources and stronger strategies for rare-cell annotation may further improve performance. The contribution of visual reasoning also varied across tissues, suggesting that future versions should determine when visual evidence is likely to be informative rather than applying it uniformly.

The skin cancer case study illustrates CellPilot not merely as an automated annotation pipeline, but as a framework for autonomous biological interpretation. Starting from raw expression profiles, the agent independently recovered coherent cellular populations, including a rare Langerhans cell population that was not explicitly represented in the provided annotation prior. Importantly, this should not be interpreted as a claim of novel biological discovery, but rather as evidence that agentic reasoning can recover biologically valid and non-obvious hypotheses beyond the analyst’s initial assumptions. The resulting annotations further supported downstream analyses of cellular communication, suggesting that the inferred cellular identities were sufficiently coherent to enable systems-level biological interpretation without manual curation.

## Methods

### CellPilot workflow formulation

CellPilot was designed to execute single-cell RNA-sequencing (scRNA-seq) analysis as a sequence of state-dependent analytical decisions rather than as a terminal cell type labeling step. The input to CellPilot is a raw or minimally processed gene expression matrix, denoted by **X** ∈ ℝ*^n^*^×*p*^, where n is the number of cells and *p* is the number of genes. Input files were provided in AnnData (`.h5ad`) or numeric matrix (`.csv`) format. A species identifier, tissue context and a marker-gene reference table could be supplied when available. The output of one CellPilot session consists of cluster-level cell type annotations, annotation support scores, intermediate analysis objects, diagnostic figures and structured execution logs.

Let *τ* = {*t*_1_, *t*_2_,…, *t_K_*} denote the set of atomic analytical tools available to the agent. These tools implement data inspection, quality control, filtering, normalization, feature selection, dimensionality reduction, neighborhood graph construction, clustering, differential expression, marker matching, label refinement, visualization and report generation. CellPilot defines a tool-constrained mapping

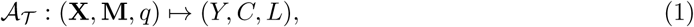

where **M** is the marker reference table, *q* is the user query or analysis instruction, 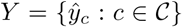 is the set of predicted labels for clusters 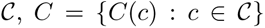 is the corresponding annotation support score, and *L* contains the structured action log, tool outputs, intermediate files and diagnostic artifacts generated during the session. All files were stored in a session-specific working directory using deterministic names.

The workflow was formulated as an iterative decision process. At iteration *t*, the agent maintains a state

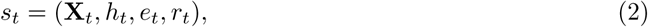

where **X***_t_* is the current AnnData representation, *h_t_* contains intermediate analytical outputs, *e_t_* records executed tool calls and returned observations, and *r_t_* records the completion status of required workflow stages. At each iteration, the controller selects one action

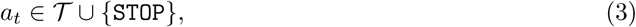

executes the corresponding tool when *a* ∊ 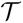, receives an observation *o_t_*, and updates the state according to

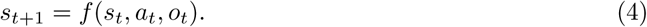

The workflow terminates when the controller emits a valid final response after completion of the required stages, or when a predefined iteration limit is reached. The required stages were data loading, quality-control assessment, filtering when required, normalization, highly variable gene selection, dimensionality reduction, neighborhood graph construction, clustering, marker detection, cell type assignment, validation and report generation.

### State-machine constrained agent orchestration

CellPilot was implemented as a state-machine controlled agent using LangGraph. The state object contained the dataset path, session directory, current AnnData file, message history, execution counter, completed workflow stages, tool-call records and returned observations. The state machine constrained the agent to select one validated analytical action at a time.

The language model controller generated actions using a restricted XML interface. A tool call was required to appear inside an `<execute>` block, and a final response was required to appear inside a `<solution>` block. Tool outputs were returned to the controller as structured `<observation>` blocks. At each iteration, the generated XML was parsed and checked before execution. Invalid XML, missing arguments or unsupported tool names were returned as structured error observations, allowing the agent to revise the next action without terminating the session.

Tool calls were executed through predefined Python wrappers with restricted arguments and session-local file paths. Each wrapper returned a structured observation containing a textual summary, output file paths when applicable, and an execution status. Runtime exceptions were captured and returned as error observations. These errors were not treated as session-level failures unless the agent reached the iteration limit or failed to produce the required final artifacts.

A self-checking step was applied before accepting a final `<solution>` block. This step checked whether all required workflow stages had been completed at least once or had been explicitly skipped with a valid reason in the action log. If required stages were missing, the final response was rejected and control returned to the agent. The self-checking step evaluated workflow completeness only; it did not use reference labels and did not judge biological correctness of the predicted annotations.

Because scRNA-seq workflows can generate long textual and visual outputs, observations were compressed before being appended to the active model context. The full observations and output files were retained in the session directory, whereas the active reasoning context retained only bounded summaries of recent observations. The maximum observation length and number of retained observations used in the experiments are reported in Supplementary Note 1.

### Language model controllers

Qwen3-8B [27] was used as the default language model controller in CellPilot. The model was deployed locally through the vLLM inference engine [47]. The controller did not directly manipulate the expression matrix. Instead, it selected from the fixed tool set described in Section 1, interpreted structured observations and decided whether additional analysis steps were required.

Each session used a fixed prompt template containing the task description, available tool schemas, XML response rules, required workflow stages and constraints on final output format. The prompt instructed the controller to inspect data properties, quality-control metrics, dimensionality-reduction results, clustering structure and marker evidence before producing the final annotation report. To reduce output variability, inference used a fixed sampling temperature of τ = 0.1 across experiments. The model name, checkpoint, serving configuration, decoding parameters and context limits are described in Supplementary Note 3.

For backbone comparison experiments, the same CellPilot workflow, tool library, prompt template and evaluation protocol were used with Qwen3-8B and GPT-4o. For prompt-only comparisons, GPTCelltype was run using marker-gene inputs without the CellPilot state machine, tool execution loop or intermediate observations. API-based models were queried with fixed prompts, and the model version and access date are reported in Supplementary Note 3.

### Atomic tool library and workflow stages

CellPilot used a library of atomic tools implemented around Scanpy and AnnData. Each tool read the current AnnData object, performed one analytical operation, wrote updated objects or figures to the session directory, and returned a structured observation to the agent. The tool library contained the following functional groups.

#### Data inspection and quality control

The input matrix was loaded into an AnnData object. Cell and gene counts, total counts per cell and the fraction of mitochondrial transcripts were computed. Mitochondrial genes were detected using species-specific prefixes. Quality-control plots were generated for gene counts, total counts and mitochondrial content. Filtering thresholds were selected by predefined quantile-based heuristics with a conservative retention rule that preserved at least 90% of cells unless the data contained clear low-quality tails. Exact default thresholds and fallback rules are described in Supplementary Note 2.

#### Normalization and feature selection

Filtered count matrices were library-size normalized and log transformed. Highly variable genes were selected using the default Scanpy procedure. If dispersion-based selection failed or returned an unstable feature set, the workflow invoked a Seurat v3-style fallback [48]. The number of highly variable genes, normalization target, minimum gene filters and fallback criteria are described in Supplementary Note 2.

#### Dimensionality reduction and graph construction

Principal component analysis was performed on the selected highly variable genes using a fixed random seed. A neighborhood graph was constructed from the top principal components, followed by UMAP projection for visualization [49]. The number of principal components, number of neighbors, distance metric and random seed are described in Supplementary Note 2.

#### Clustering

Graph-based clustering was performed using the Leiden algorithm [50]. Cluster labels were stored in the AnnData object and used for downstream differential expression and annotation. The default resolution and any agent-selected resolution adjustments were recorded in the session log. The agent was allowed to repeat clustering with a modified resolution when the returned observations indicated severe over-clustering, under-clustering or poor separation in the diagnostic plots.

#### Differential expression and marker extraction

For each cluster, differentially expressed genes were computed against the remaining cells using rank-based testing as implemented in Scanpy, consistent with established practice in single-cell differential expression analysis [51, 52]. The top-ranked genes for each cluster were written to the AnnData object and to a session-level marker table. These cluster markers were used for marker-database matching and, when needed, language-model assisted label refinement. The differential expression test, ranking statistic and number of retained genes are described in Supplementary Note 2.

### Marker-based annotation and support scoring

Let 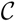 denote the set of clusters produced by CellPilot and let 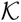 denote the set of candidate cell types in the marker reference. For each cluster *c* ∈ 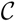 and candidate cell type *k* ∈ 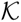, a marker-hit count *h*(*c*, *k*) was computed by intersecting the top differentially expressed genes of cluster c with the marker set of cell type *k*. The marker-matched label was assigned as

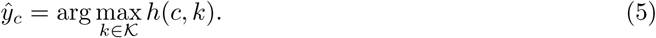

If 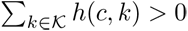, the marker-derived support score was defined as

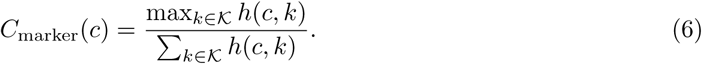

Clusters with no marker hits were marked as unassigned after marker matching.

Unassigned clusters were passed to a label-refinement prompt. The prompt contained the cluster identifier, top differentially expressed genes, available marker overlaps, tissue context and cluster metadata. The model returned a JSON object containing a predicted cell type, a support score in [0, 1] and a short justification based on marker evidence. The response was accepted only if it conformed to the predefined JSON schema. Invalid responses were assigned an unassigned label with a conservative support score.

The final support score for cluster c was

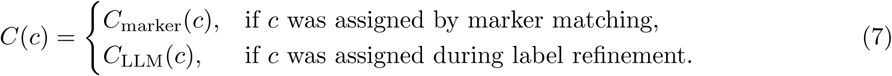

The support score was used as a triage measure for prioritizing annotations for review. It was not treated as a calibrated posterior probability of correctness.

### Visual observation enrichment

CellPilot optionally used a vision-language module to convert diagnostic figures into textual observations. Vision-language models combine large-language-model reasoning with visual encoders trained to interpret images alongside textual instructions [53, 54]. The module was applied to figures generated during the workflow, including quality-control violin plots, principal component variance plots, UMAP embeddings, marker dot plots and feature plots. In the experiments using visual observations, Qwen3-8B-VL-Thinking was used as the visual summarization model.

For each figure, the module generated a structured textual summary of salient visual patterns. Quality-control summaries described distributional features such as extreme tails or high mitochondrial content. Embedding summaries described cluster separation and visually ambiguous regions. Marker-expression summaries described cluster specificity and co-expression patterns. These summaries were appended to the agent state as auxiliary observations. The VLM module did not assign cell type labels directly and did not modify numerical tool outputs.

Figure summaries were cached by file path and metadata to avoid repeated processing of the same image. If visual summarization failed or returned invalid text, an empty observation was recorded and the workflow continued. Ablation experiments disabled this module while keeping all other tool calls, prompts and evaluation procedures unchanged.

### Benchmark datasets

CellPilot was evaluated on three publicly available scRNA-seq benchmarks spanning human and mouse tissues. For CellPilot, each dataset was analyzed from the raw count matrix or the count layer provided by the original resource. No manual preprocessing, manual cluster selection or post hoc correction of CellPilot annotations was applied before evaluation.

a. *GTEx* [37]: a human single-cell and single-nucleus benchmark containing seven tissues with curated cell type annotations. Raw count matrices were used as input.
b. *Tabula Sapiens* [38]: a multi-organ human cell atlas containing twenty-four tissues with harmonized reference labels. Raw count matrices were analyzed directly.
c. *Mouse Cell Atlas* [39]: a mouse cross-tissue atlas with manually curated cell type labels. Raw count matrices were used as provided by the source dataset.

Reference labels from the original studies were used only for evaluation. They were not supplied to CellPilot during workflow execution. Dataset-level statistics, including the number of cells, tissues, reference clusters and reference cell type labels, are summarized in Supplementary Note 5.

### Baseline methods and input assumptions

CellPilot was compared with representative reference-based, marker-based and LLM-based annotation methods. Because these methods use different input assumptions, we separated method inputs from evaluation units. Cluster-dependent baselines were evaluated under their intended pre-defined cluster settings, whereas CellPilot performed de novo preprocessing, clustering, marker discovery and annotation from raw count matrices. All predictions were subsequently evaluated against the same dataset-defined reference annotations.

a. *SingleR* [13]: SingleR was run as a reference-based cell-level annotation method. It assigned labels by comparing query profiles with reference expression signatures using correlation-based scoring. Cell-level predictions were aggregated to the reference-cluster level by majority vote.
b. *ScType* [12]: ScType was run as a marker-based cluster annotation method using curated marker gene sets. For datasets where tissue-specific marker references were available, ScType was applied to the dataset-provided cluster structure.
c. *GPTCelltype4* [23]: GPTCelltype4 was run as a prompt-based LLM annotation baseline using differential gene lists from pre-defined clusters. The marker size parameter was set to 10 following the original protocol, and tissue context was supplied according to the dataset.
d. *CASSIA* [24]: CASSIA was run as a structured LLM-based annotation baseline using marker genes and tissue context. Default parameters were used unless otherwise specified by the original implementation.

No manual correction, relabeling or post hoc filtering was applied to baseline outputs. The input data, cluster assumptions, marker sources, model versions and runtime settings for each method are summarized in Supplementary Note 5.

### Evaluation alignment for de novo and reference clusters

CellPilot produces labels on de novo clusters, whereas several baselines operate on dataset-defined reference clusters. To evaluate all methods on a common unit, we projected method outputs onto the reference clusters provided by each benchmark dataset.

For CellPilot, every cell inherited the predicted label of its CellPilot cluster. These cell-level labels were then grouped by the dataset-defined reference cluster. Two aggregation schemes were used. In the reference-cluster vote scheme, the modal CellPilot label among cells in a reference cluster was selected as the predicted label for that reference cluster. In the reference-cluster mean-score scheme, each cell first received an ontology score according to the label of its CellPilot cluster, and scores were then averaged within each reference cluster. For baselines that directly returned labels on reference clusters, the returned cluster label was scored directly. For SingleR, cell-level predictions were aggregated within each reference cluster by majority vote before scoring.

This alignment used reference clusters only for evaluation. CellPilot did not use reference clusters, reference marker lists or reference labels during workflow execution.

### Ontology-based categorical scoring

Annotation performance was evaluated using the hierarchical scoring protocol of Hou and Ji [23], which compares predicted and reference labels through the structure of the Cell Ontology [55]. Let *ŷ_r_* denote the predicted label for reference cluster *r*, and let 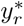 denote the corresponding reference label. A categorical score *S*(*r*) ∈ {0, 0.5, 1} was assigned as follows:

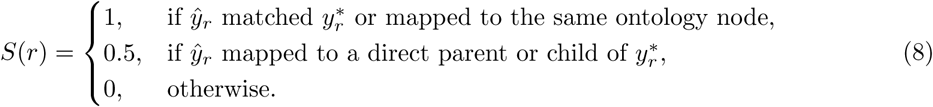

For labels with different annotation granularity, a broader parent label was counted as partially correct when it corresponded to the immediate ontology parent of the reference label. Manual mappings and unresolved terms were recorded before scoring and applied consistently across all methods.

For a tissue with reference clusters 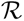, the tissue-level score was computed as

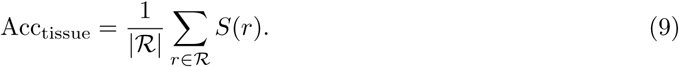

Dataset-level scores were computed by averaging tissue-level scores. Unless otherwise specified, statistical comparisons across methods were performed on matched tissue-level scores using two-sided paired Wilcoxon signed-rank tests. When a dataset yielded only a single dataset-level value for a comparison, no inferential test was reported for that dataset alone.

### Semantic similarity analysis

Semantic similarity between predicted and reference cell type names was computed as a complementary analysis for non-exact annotations. Cell type terms were embedded using SapBERT [56]. For a predicted term *ŷ_r_* and reference term 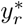, cosine similarity was computed as

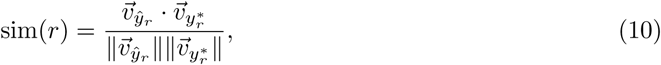

where 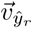 and 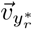 are the SapBERT embeddings of the predicted and reference terms. Semantic similarity was computed for predictions with *S*(*r*) = 0 or *S*(*r*) = 0.5 to characterize the biological proximity of imperfect predictions. Exact matches were excluded from this analysis.

### Cell-level and cluster-level stratified analyses

For cell-level summaries, each cell was assigned the score of the reference cluster to which it belonged after prediction alignment. Cells were then grouped by dataset, tissue, broad cell category, reference-label granularity and cluster size. Cluster size was defined using the number of cells in the dataset-defined reference cluster. Small, medium and large clusters were defined according to the thresholds defined in Supplementary Note 7. Full-match, partial-match and mismatch proportions were computed within each group.

Broad cell categories were generated by mapping fine-grained reference labels to manually curated parent categories. These mappings were used only for stratified analysis and visualization, not for primary scoring. The broad-category mapping is described in Supplementary Note 7.

### Support-score assessment

The relationship between CellPilot support scores and annotation correctness was evaluated after reference-cluster alignment. For each tissue, mean predicted support was computed across reference clusters, and compared with the corresponding tissue-level ontology score. Pearson correlation coefficients were reported as descriptive measures. Support-score threshold analysis was performed by retaining predictions with *C*(*c*) above a threshold and computing the full-match rate and retained-cluster fraction at each threshold. These analyses were used to evaluate whether support scores could prioritize annotations for review. They were not used to treat support scores as calibrated probabilities.

### Workflow reliability analysis

Each tool call was logged with the tool name, arguments, execution status, returned observation, generated files and runtime messages. A tool call was considered successful if it completed without an uncaught exception and returned the expected structured output. The tool success rate was computed as

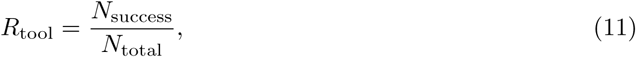

where *N*_success_ is the number of successful tool calls and *N*_total_ is the total number of attempted tool calls.

A session was considered complete if it produced a final annotation table, an updated AnnData object, diagnostic visualizations and an annotation report after executing or explicitly skipping all required workflow stages. The workflow completion rate was computed as

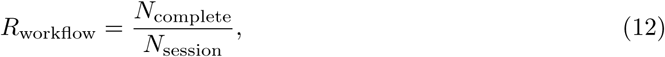

where *N*_complete_ is the number of complete sessions and *N*_session_ is the total number of sessions.

Per-tool failure rates were computed by grouping tools into data inspection, preprocessing, dimensionality reduction, clustering, marker detection, marker query, annotation refinement, visualization and reporting. When a tool failed, the error observation was retained in the action log and made available to the agent in the next iteration. The number of tool calls per session and the frequency of validation-triggered refinement were recorded to characterize execution behavior.

### Implementation and reproducibility

CellPilot was implemented in Python 3.11. Single-cell preprocessing, dimensionality reduction, clustering, differential expression and visualization were implemented using Scanpy 1.10 and AnnData. Agent orchestration was implemented using LangGraph. Qwen3-8B was served locally using vLLM. The optional visual summarization module used Qwen3-8B-VL-Thinking. Package versions, model checkpoints, decoding parameters and hardware configuration are described in Supplementary Notes 1 and 3.

All analyses used fixed random seeds for PCA, neighborhood graph construction, UMAP and Leiden clustering when the underlying implementation supported seeded execution. Each session was executed in a dedicated working directory. Intermediate AnnData files, diagnostic plots, marker tables, annotation tables, action logs and final reports were saved using deterministic filenames. Prompt templates, tool schemas, marker reference files and evaluation scripts were versioned with the codebase.

The marker database used by CellPilot was stored as a versioned reference file. The database version, source, download date, supported species and number of included cell type entries are described in Supplementary Note 6. API-based baselines were run with the model versions and access dates listed in Supplementary Note 3.

## Declarations

### Data availability

All datasets used in this study are publicly available and were obtained without restriction. The GTEx single-nucleus cross-tissue reference [37] was downloaded from the Single Cell Portal (study SCP1479) and the GTEx Portal (https://gtexportal.org). The Tabula Sapiens multi-organ human atlas [38] was obtained from the Tabula Sapiens data portal (https://tabula-sapiens-portal.ds.czbiohub.org) and the CELLxGENE collection. The Mouse Cell Atlas [39] was retrieved from the Gene Expression Omnibus under accession GSE108097 and the associated Figshare repository. For each dataset, raw count matrices and the original authors’ curated cell-type annotations were used as input and as ground-truth labels, respectively.

### Author contribution

S.J. and Z.W. conceived and designed the study. S.J. conducted the experiments and wrote the manuscript. C.Q. conducted the experiments and contributed to manuscript writing. X.S. conducted partial experiments. Y.C. analyzed the experimental results and was responsible for biological validation. Z.W. supervised the overall study.

## Supplementary Information

This Supplementary Information accompanies the manuscript entitled *CellPilot: an agentic framework that pilots small language models through autonomous single-cell annotation*. It provides implementation-level, methodological, and evaluation details that complement the main text and support reproducibility, technical transparency, and independent assessment of the framework. Specifically, the Supplementary Information documents the agent system architecture and implementation details, complete tool function specifications, prompt engineering and chain-of-thought design, and the integration of the vision-language model module. It further describes the benchmark design, dataset preparation procedures, and fairness controls used for comparative evaluation, as well as the construction and use of the marker database, annotation provenance tracking, and the hierarchical evaluation rules applied in scoring prediction outcomes. Finally, it presents additional analyses of cell-level annotation performance across annotation granularity, broad cell type categories, cluster size, and tissue context. Together, these materials are intended to provide a complete technical record of the CellPilot framework beyond the space constraints of the main manuscript.

### Supplementary Note 1: Agent System Architecture and Implementation Details

This note provides implementation-level details that supplement the architectural overview and algorithmic description presented in Section 4.2 of the main text. The focus is on engineering decisions, specific parameter values, and subsystem designs not covered in the main Methods.

#### 1.1 State Fields Beyond the Core Architecture

The `AgentState` object described in Section 4.2 is implemented as a Python `TypedDict` containing seven fields. In addition to the five fields noted in the main text (user query, current dataset path, session-specific working directory, message history, execution counter, and tools-executed record), two additional fields support workflow control: `plan_generated`, a boolean flag that distinguishes the initial planning turn from subsequent execution turns to prevent the routing logic from misinterpreting a plan-only response as a stalled state; and `session_id`, a UUID-based identifier used to construct the session directory path and to tag logs for multi-session deployments.

#### 1.2 LangGraph Implementation and Routing Details

The state-machine control loop described in Section 4.2 and Algorithm 1 is compiled via LangGraph’s `StateGraph` API into a directed graph with conditional edges. Two routing details not captured by Algorithm 1 are relevant for reproducibility. First, the `execute` node is connected back to `generate` via a fixed unconditional edge rather than a conditional one, which guarantees that every tool invocation is followed by a new LLM reasoning step. There is no execution path in which two consecutive tool calls occur without an intervening reasoning iteration. Second, the self-critic node is inserted between the `generate` node and the terminal `END` node only when the critic mechanism is enabled. When disabled, the `<solution>` directive transitions directly to `END`, reducing latency by one additional LLM call per session.

#### 1.3 Execution Sandboxing and Path Resolution

Tool calls are executed within a sandboxed environment managed by the `ExecutionEngine` class, a subsystem not described in the main text. Before each execution, the working directory is changed to the session-specific directory via `os.chdir`, ensuring that all relative file paths resolve within the session scope. Upon completion or failure, the original working directory is restored. File paths appearing in generated code are automatically rewritten to absolute paths using a regex-based resolver that checks for the existence of candidate files first in the session directory and then in the global working directory. This path rewriting prevents file-not-found errors that would otherwise arise when the language model generates code referencing filenames without full paths.

Tool outputs are captured by redirecting `stdout` and `stderr` into string buffers, intercepting any generated image files detected either through regex matching of file extensions in output text or through explicit `matplotlib.Figure` objects in the execution namespace, copying images into a `figures/` subdirectory under the session directory with UUID-prefixed filenames to avoid collisions, and returning a standardized result dictionary containing an `ok` boolean, the captured output text, a list of image records, and an updated data path if the tool modified the AnnData object.

#### 1.4 Session Directory Structure and Artifact Persistence

The main text notes that each session operates within a dedicated working directory with deterministic naming conventions (Section 4.10). Here we specify the complete directory layout. Each session is created under a global sessions root (`sc analysis/sessions/<uuid>/`). The uploaded dataset is copied into this directory at session initialization to ensure immutability of the original file. All intermediate AnnData objects are saved with the following deterministic suffixes:

- `<dataset>.qc.h5ad` — after QC metric calculation
- `<dataset>.qc.filtered.h5ad` — after cell filtering
- `<dataset>.norm.h5ad` — after normalization and log-transformation
- `<dataset>.hvg.h5ad` — after highly variable gene selection
- `<dataset>.norm.hvg.norm.scaled.h5ad` — after scaling of HVG-subset data
- `<dataset>.pca.h5ad` — after PCA computation
- `<dataset>.neighbors.h5ad` — after neighbor graph construction
- `<dataset>.umap.h5ad` — after UMAP embedding
- `<dataset>.leiden r`*X*`.h5ad` — after Leiden clustering at resolution *X*
- `<dataset>.deg.h5ad` — after differential expression analysis
- `<dataset>.annotated.h5ad` — final annotated object

In addition, the session directory stores the following log and diagnostic files:

- `figures/` — all generated visualizations, with UUID-prefixed filenames
- `debug llm requests.log` — abbreviated LLM request previews (last 5 messages per call)
- `debug llm requests full.log` — complete, untruncated LLM request history for each call
- `debug llm responses.log` — all raw LLM responses
- `execution outputs.log` — tool execution outputs and observation text
- `qwen reasoning.log` — full reasoning traces when Qwen thinking mode is enabled
- `timing.json` — overall session runtime (start time, end time, elapsed seconds, total tool time)
- `timing records.json` — per-step timing breakdown with timestamps

This logging structure enables post hoc auditing of the analytical trajectory followed in each session.

#### 1.5 Context Window Management: Parameter Specifications

Section 4.2 of the main text describes the general context management strategy, including observation truncation and bounded retention of recent observations. Here we specify the three-level truncation hierarchy and its default parameter values as implemented.

At the *observation level*, individual tool outputs undergo two sequential truncations: first by line count (default: 150 lines retained from the head, plus the last 5 lines with an inserted skip marker indicating the number of omitted lines), and subsequently by character count (default: 2,000 characters, with a trailing truncation marker). This dual truncation preserves key summary statistics at the beginning of tool outputs and tail-end status information while discarding verbose intermediate content.

At the *history level*, a category-aware pruning method operates before each LLM invocation, processing the message history in reverse chronological order and independently limiting three message categories: observation messages (those containing `<observation>` tags, capped at the 6 most recent), AI reasoning messages (capped at the 8 most recent), and user messages (capped at the 3 most recent turns). The two most recent messages are always retained regardless of category limits. The total message count is bounded at 40 messages. All parameters are configurable at agent initialization and can be adjusted for language models with different context window sizes.

At the *VLM summary level*, image summaries are constrained to 100 words each, enforced by post hoc word-count truncation even if the VLM exceeds the prompt instruction, with at most 2 image summaries generated per tool invocation.

#### 1.6 Error Handling: Specific Safeguards

Section 4.2 describes the general error recovery strategy in which failures are returned as structured observations that allow the model to revise its plan. Here we document three additional safeguards implemented in the code.

First, a global recursion limit of 100 LangGraph iterations is set via the `recursion_limit` configuration parameter. This prevents infinite cycling in edge cases where the model repeatedly generates unparseable or non-progressing outputs.

Second, a secondary safety check within the `generate` node forces session termination with an automatic `<solution>` message if the execution counter exceeds 50 tool invocations, providing a tighter upper bound than the graph-level recursion limit.

Third, when a `<solution>` tag is emitted, the agent verifies that all required tools have been executed by checking the `tools_executed` list against a required set that currently includes `tool_validate_annotation`. Because this validation step internally invokes `tool_generate_annotation report`, the final quality check and report generation are coupled. If the requirement is not satisfied, the proposed solution is intercepted, a reminder is injected into the conversation history, and control returns to `generate`.

#### 1.7 Custom Marker File Injection

When a user provides a custom marker gene database, the file is saved to the session directory and its path is recorded in a `marker_file_path.txt` sentinel file. During execution, if the agent-generated code calls `tool_query_markers` without specifying a `marker_csv_path` argument, or explicitly sets `marker_csv_path=None`, the execution engine automatically injects the custom marker file path into the function call before execution. This mechanism allows the language model to generate generic, path-agnostic tool calls while ensuring that user-provided marker databases are consistently applied across the session.

#### 1.8 Algorithm of State-machine control loop used by CellPilot

##### Algorithm 1

**State-machine control loop used by CellPilot**

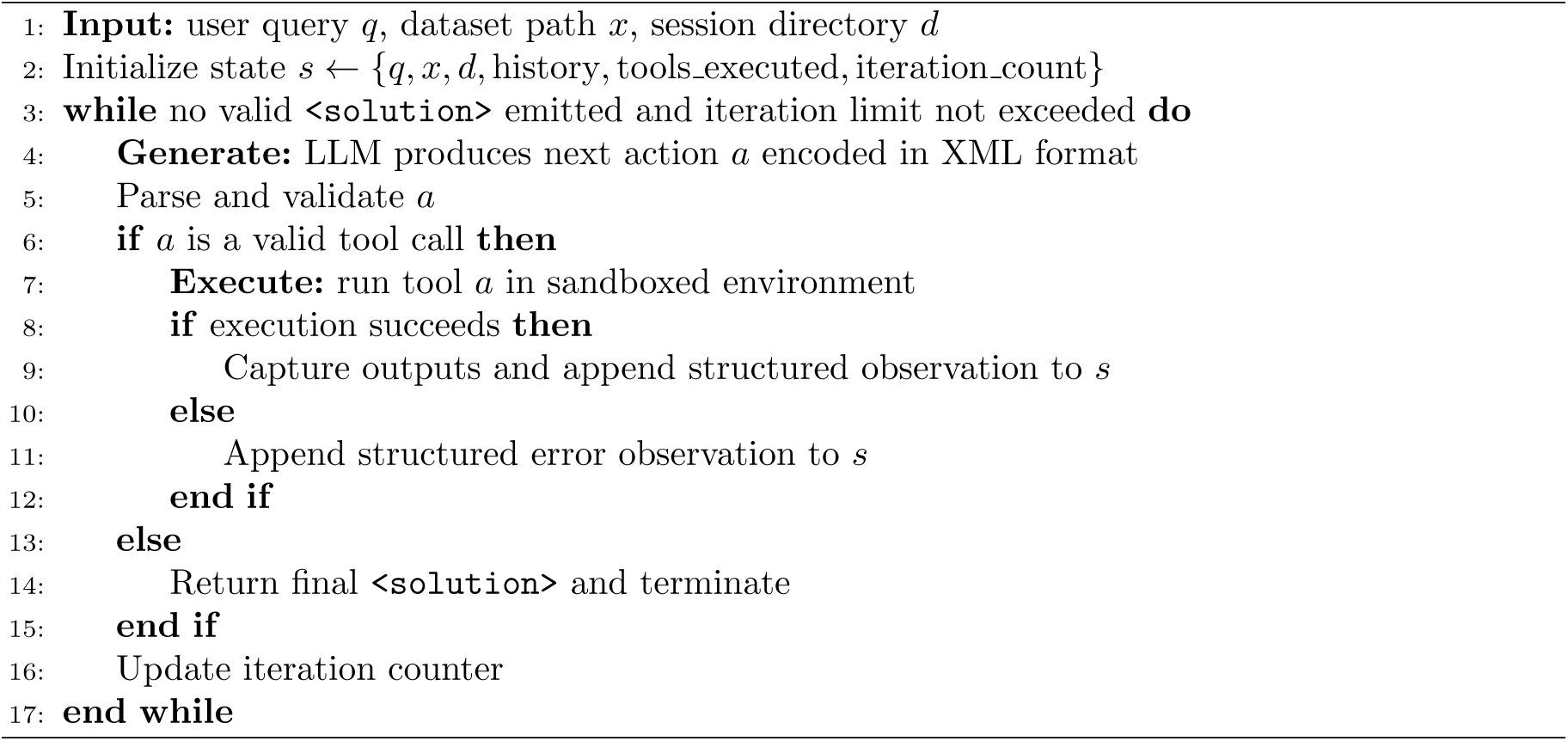

### Supplementary Note 2: Tool Function Specifications

This note documents the complete function signatures, default parameter values, and implementation-specific behaviors of the 16 tools in the CellPilot toolkit. Section 4.5 of the main text describes the analytical logic and mathematical formulations underlying the preprocessing, clustering, and annotation stages. Here we focus on software interfaces and operational behavior relevant to reproduction and extension of the toolkit.

All tools accept the current AnnData file path as their primary argument, operate on the shared AnnData object, persist updated objects and generated visualizations to the session directory, and return a structured ToolResult containing an execution status flag, a textual summary, and references to any output files.

#### 2.1 Data Ingestion and Inspection

**`tool_inspect_data(file path)`** accepts `.h5ad` and `.csv` input formats. For AnnData files, it extracts matrix dimensions, available annotation fields in `obs` and `var`, and summary statistics of the expression values. A heuristic classifier determines whether the data appear to represent raw counts, log-transformed values, or scaled values by examining the value range, checking for the presence of negative entries, and inspecting the distribution shape. An expression histogram is saved to the session directory to allow visual confirmation of the detected preprocessing state. This step informs the agent’s initial planning decision by indicating whether normalization and transformation should be executed or skipped.

#### 2.2 Quality Control

**tool calculate qc metrics(file path)** wraps Scanpy’s `sc.pp.calculate qc metrics` and automatically detects mitochondrial genes by searching gene name prefixes `MT-`, `mt-`, and `Mt-` to accommodate human, mouse, and mixed-case conventions. The tool computes per-cell total counts, number of genes detected, and mitochondrial count percentage, and reports quantile distributions at the 5th, 25th, 50th, 75th, and 95th percentiles for each metric. A multi-panel violin plot is generated alongside the numeric output.

**tool filter by qc(file path, min genes=200, max genes=None, max mito=20.0)** applies cell-level quality filters with configurable thresholds. The default values are intentionally conservative: `min genes=200` removes only cells with extremely low gene detection, `max genes` is left unset by default to avoid removing large cells without evidence of doublet contamination, and `max_mito=20.0` accommodates tissues with moderate mitochondrial content. These defaults are designed to satisfy the `≥`90% cell retention guideline described in the main text. The tool reports the number of cells before and after filtering and the retention percentage.

#### 2.3 Normalization and Feature Selection

**tool normalize data(file path, normalize=True, log transform=True, scale=False, target sum=10000)** provides fine-grained control over three distinct preprocessing operations through independent boolean flags. Library-size normalization scales each cell to a target sum of 10,000 counts by default. Log transformation applies `log1p`. Scaling centers each gene to zero mean and clips values at ±10 to limit the influence of outlier expression values. Importantly, the `scale` flag defaults to `False` because scaling must occur after highly variable gene selection to preserve variance information. The tool enforces this order internally and raises a warning if scaling is requested on data that have not yet undergone HVG selection.

**tool hvg(file path, n top genes=2000, flavor=‘seurat v3’)** identifies highly variable genes using Scanpy’s `sc.pp.highly_variable_genes`. The `seurat_v3` flavor operates on raw count data and is preferred when available. However, if the input matrix contains non-integer values, as occurs after log transformation, the tool automatically falls back to the `seurat` flavor, which is compatible with transformed data. Selected genes are flagged in `adata.var[‘highly variable’]`, and the AnnData object is saved with a `.hvg` suffix.

#### 2.4 Dimensionality Reduction

**tool run pca(file path, n comps=50)** computes principal component analysis on the HVG-filtered expression matrix. A variance-explained elbow plot is generated automatically and saved to the session directory. PCA embeddings are stored in `adata.obsm[‘X pca’]` and the per-component variance ratios in `adata.uns[‘pca’][‘variance ratio’]`.

**tool run neighbors(file path, n neighbors=15, n pcs=30)** constructs a k-nearest neighbor graph from the top principal components. The resulting connectivities and distances matrices are stored in adata.obsp.

**tool run umap(file path, random state=0)** computes two-dimensional UMAP coordinates for visualization. The fixed random seed ensures reproducibility of visual outputs for identical inputs.

#### 2.5 Clustering and Differential Expression

**tool run leiden cluster(file path, resolution=0.8, random state=0)** performs Leiden community detection on the precomputed neighbor graph. Cluster assignments are stored in both `adata.obs[‘leiden’]` and a standardized `adata.obs[‘cluster’]` column, the latter serving as the canonical cluster identifier for downstream annotation tools.

**tool visualize clusters(file path)** generates a UMAP scatter plot colored by cluster assignments and saves the figure as a PNG file. This visualization can be consumed by the VLM module (Supplementary Note 4) to produce a structured assessment of cluster separation quality.

**tool find deg(file path, n genes=25, method=‘wilcoxon’)** identifies differentially expressed genes per cluster using Scanpy’s `sc.tl.rank genes groups` with the Wilcoxon rank-sum test as the default method. For each cluster, the top-ranked genes are returned along with their scores, adjusted p-values, and log fold changes.

#### 2.6 Annotation

**tool query markers(file path, marker file=None, top k=50)** cross-references the top `K` differentially expressed genes per cluster against a marker gene database. When `marker_file` is not provided, the tool uses the built-in default database; when a custom marker file is supplied by the user, it is automatically injected by the execution engine (Supplementary Note 1, Section 1.7). For each cluster, the function computes marker hit counts for all candidate cell types and assigns the predicted label and confidence score as defined in Supplementary Note 6. Predictions and confidence values are written to `adata.obs[‘predicted cell type’]` and `adata.obs[‘predicted confidence’]`. Clusters with zero total marker hits are labeled as `Unassigned` and flagged for subsequent LLM-based refinement.

**tool validate annotation(file path)** performs post-annotation quality checks, including verification of the total number of clusters, the cluster size distribution, the count of unassigned clusters, and the number of clusters with confidence scores below 0.5. Warnings are emitted as structured text in the tool output but do not halt the workflow. This tool also internally invokes `tool_generate_annotation report`.

**tool llm annotate unassigned(file path)** is invoked only when unassigned clusters remain after marker-based annotation. For each such cluster, the tool constructs a structured prompt containing the top differentially expressed genes, any partial marker matches, cluster size, and mean QC statistics. The language model is instructed to return a JSON object with three fields: `cell type`, `confidence`, and `reasoning`. Responses that do not conform to this schema are assigned a default confidence of 0.3. Refined labels and associated confidence values are stored in `adata.obs[‘llm annotate’]` and `adata.obs[‘llm confidence’]`, respectively.

#### 2.7 Reporting

**tool generate annotation report(file path)** produces a multi-panel summary figure that consolidates the key outputs of the entire workflow into a single visualization for downstream auditing. The figure includes a UMAP embedding colored by cluster identity, a UMAP colored by predicted cell type, per-cell overlays of QC metrics, and marker gene expression patterns across annotated populations. This report is saved as a high-resolution PNG file in the session directory.

### Supplementary Note 3: Prompt Engineering and Chain-of-Thought Design

This note presents the complete prompt templates used by CellPilot, including the system prompt that governs the agent’s reasoning behavior, the structured XML protocol for tool invocation, and the auxiliary prompts for self-critique and vision-language model integration. Section 4.2 of the main text describes the role of these prompts within the state-machine architecture. Here we provide the verbatim prompt text to support reproducibility and independent evaluation of the prompt design.

### 3.1 System Prompt

The system prompt is constructed at agent initialization and remains fixed throughout the session. It comprises six functional sections: role specification and output format constraints, preprocessing order enforcement, quality control filtering guidelines, clustering parameter recommendations, workflow and execution rules, and the dynamically generated tool list. The complete prompt is shown in Box 1.

The prompt employs several design strategies that are intended to improve instruction adherence in compact reasoning models. Domain-specific constraints are phrased as imperative rules rather than suggestions. Correct usage examples are included explicitly. The preprocessing order is reinforced through both a numbered sequence and a worked code example. QC filtering guidance is framed as quantile-based heuristics with concrete numerical examples, converting a qualitative judgment into an operational decision rule.

The tool list section is generated dynamically from the registered tool functions using Python’s `inspect.signature`, ensuring that the prompt reflects the exact function signatures available in the current toolkit version.

#### Box 1.

**System Prompt (verbatim, tool list abbreviated)**

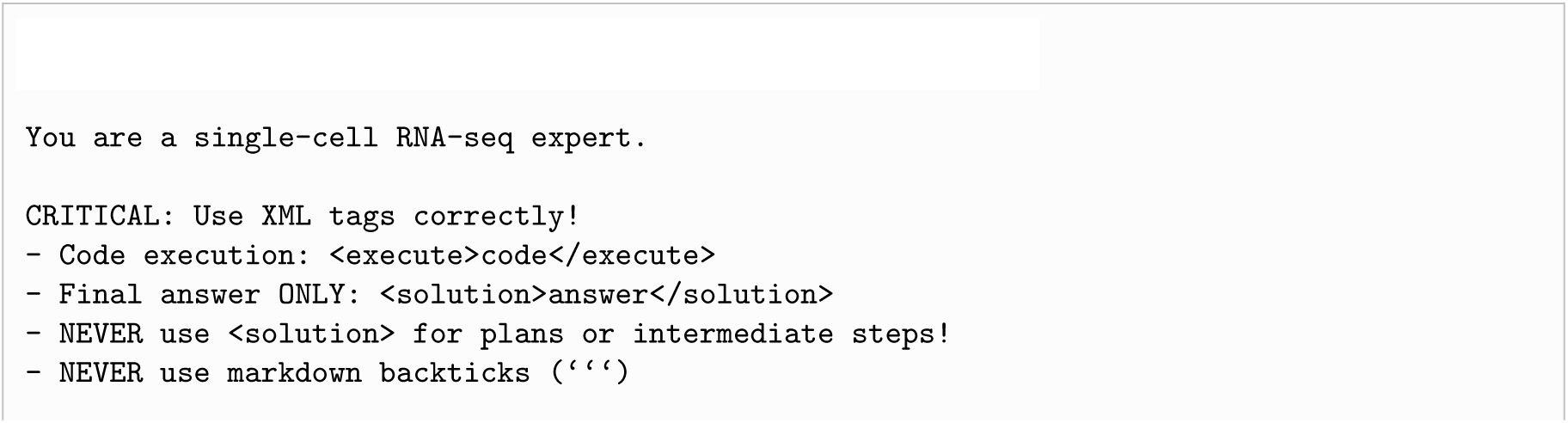
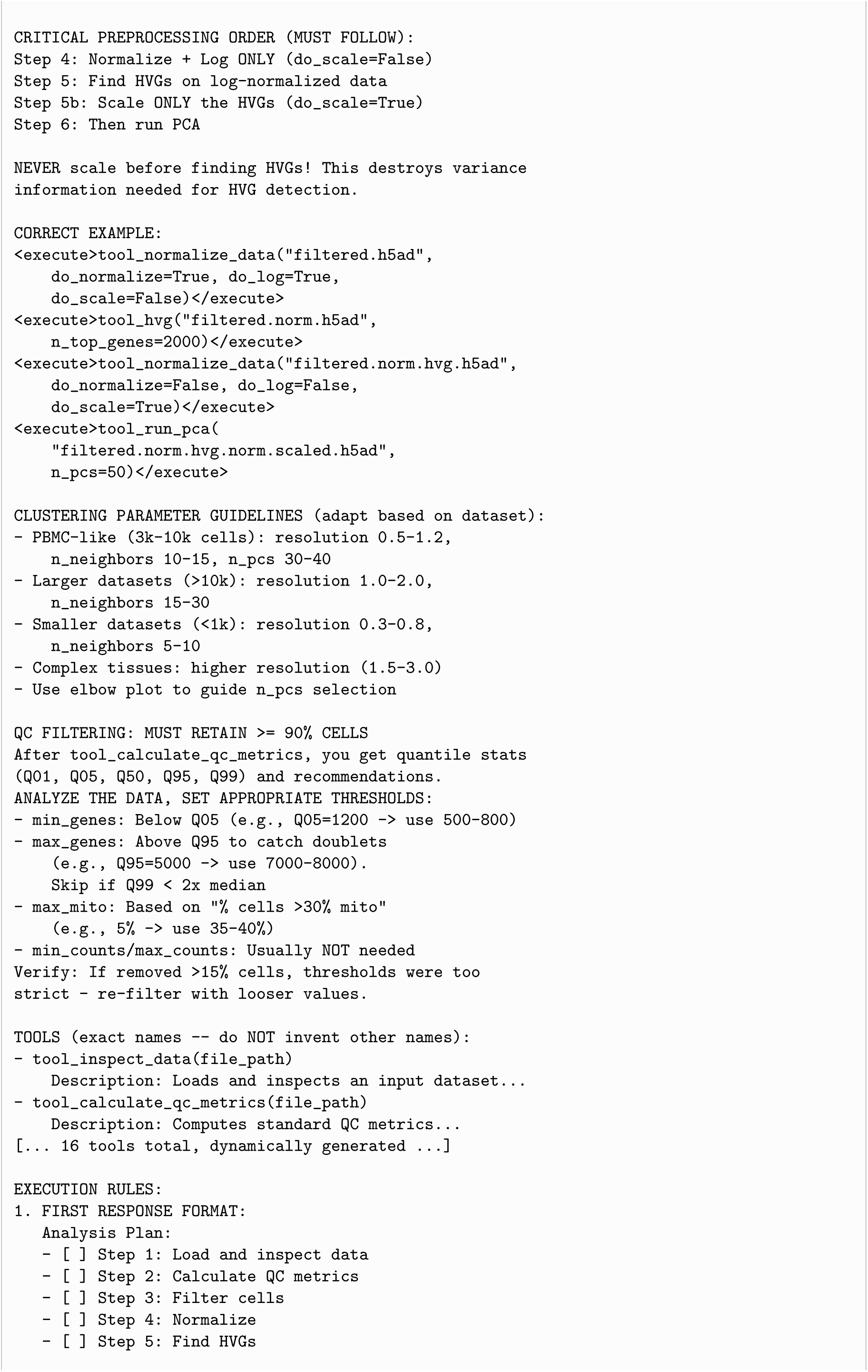
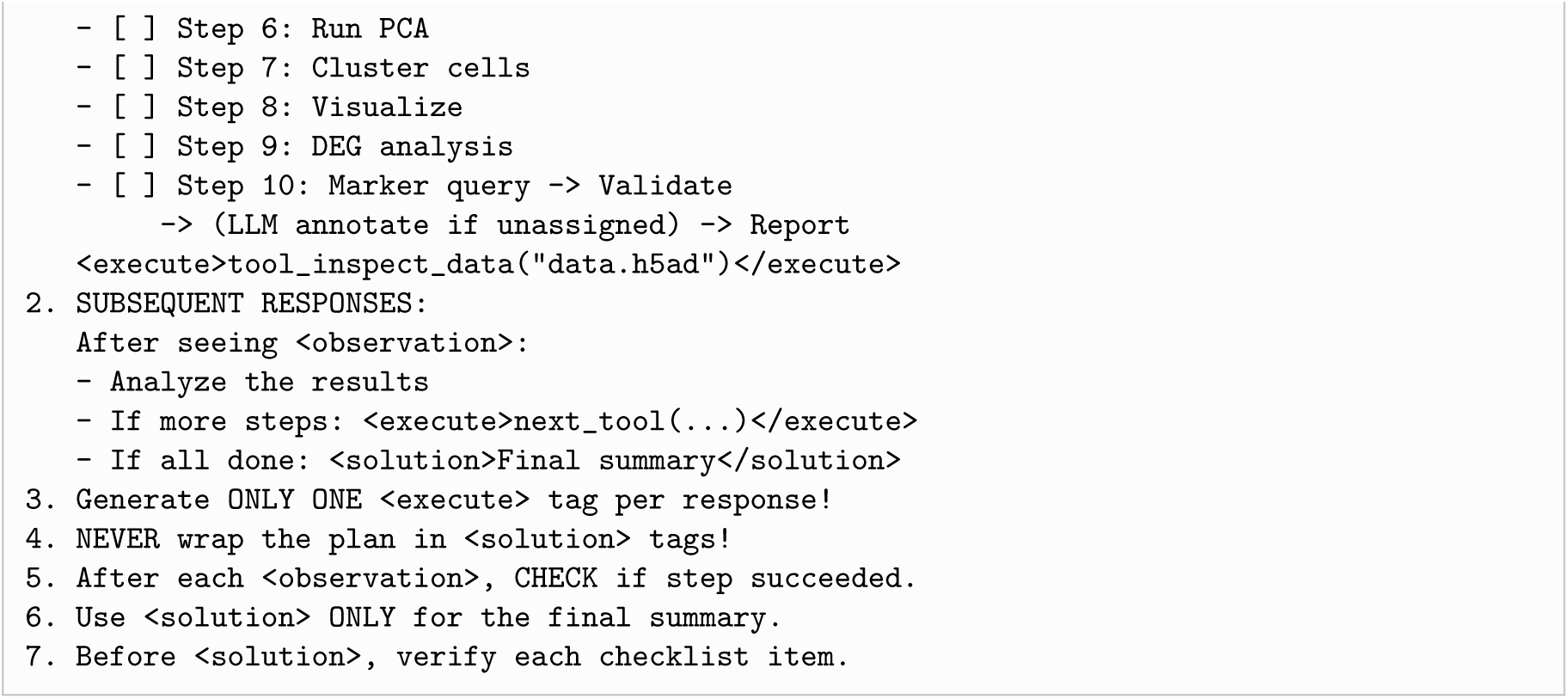

### 3.2 Dynamic Prompt Compression

CellPilot supports an alternative *dynamic* prompt mode designed for models with smaller context windows. In this mode, rather than including the full system prompt at every LLM invocation, the system selects a subset of tools relevant to the current workflow stage and constructs a shorter prompt containing only the essential formatting rules and the selected tool signatures. Tool selection operates by maintaining the canonical execution sequence and including a sliding window of up to six tools centered on the next unexecuted tool, supplemented by a fixed set of core tools (`tool_inspect_data`, `tool_normalize_data`, `tool_visualize_clusters`, `tool_validate_annotation`, `tool_generate_annotation_report`) that are always available. The compressed prompt retains the critical formatting constraints and preprocessing order but omits the more detailed QC guidelines, clustering parameter recommendations, and usage examples.

### 3.3 Structured XML Protocol and Interaction Examples

The XML protocol governs all communication between the language model and the execution engine. Three tag types are defined: `<execute>` encloses a single tool call to be executed, `<observation>` wraps the structured output returned after execution, and `<solution>` marks the final analytical summary. The agent is constrained to emit exactly one `<execute>` block per response, enforcing atomic tool execution. This constraint simplifies error isolation because a failed tool call yields a single, unambiguous observation.

Box 2 illustrates a representative three-turn interaction excerpt showing the XML protocol in practice.

#### Box 2.

**Representative interaction excerpt (XML protocol)**

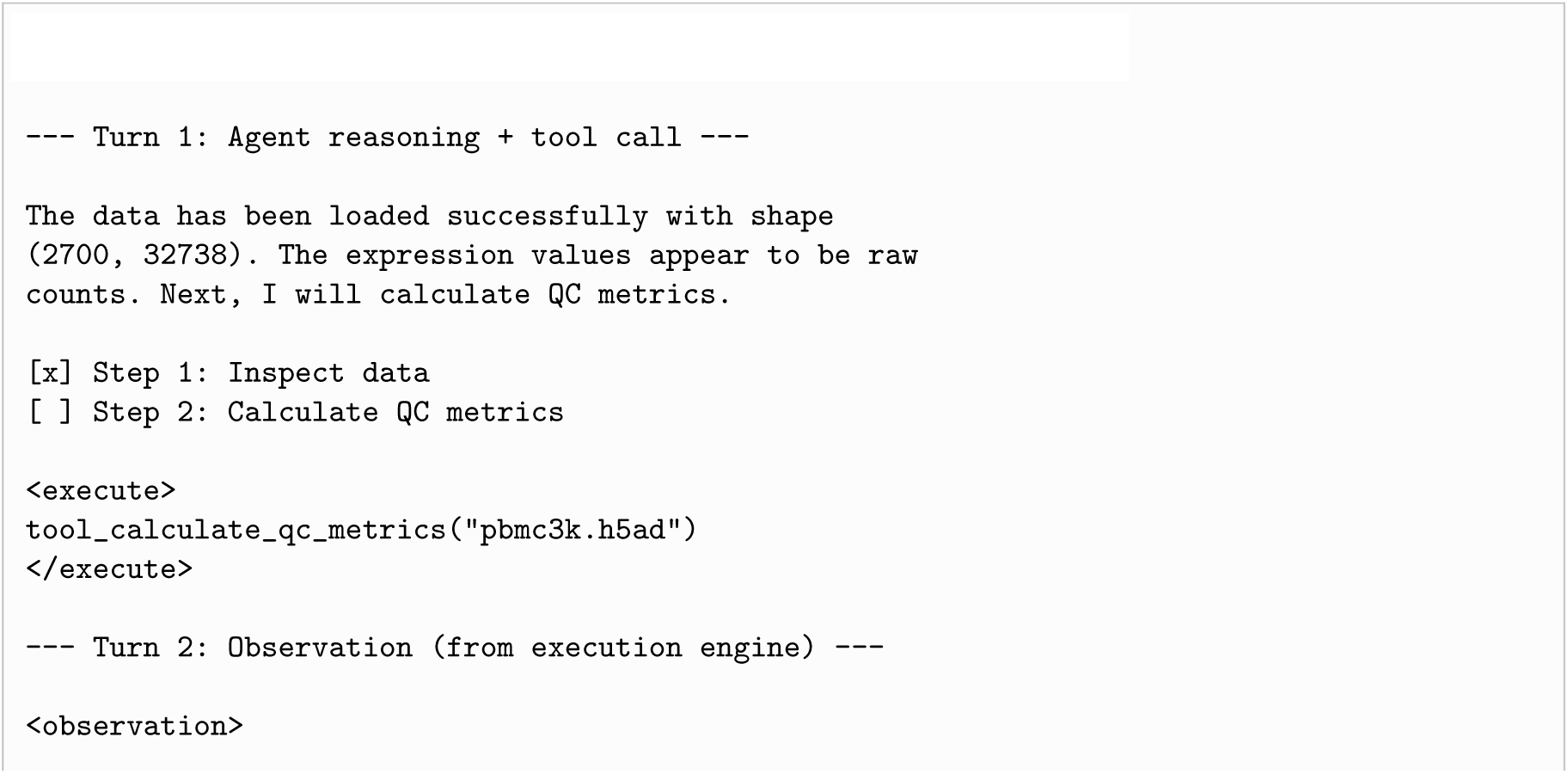
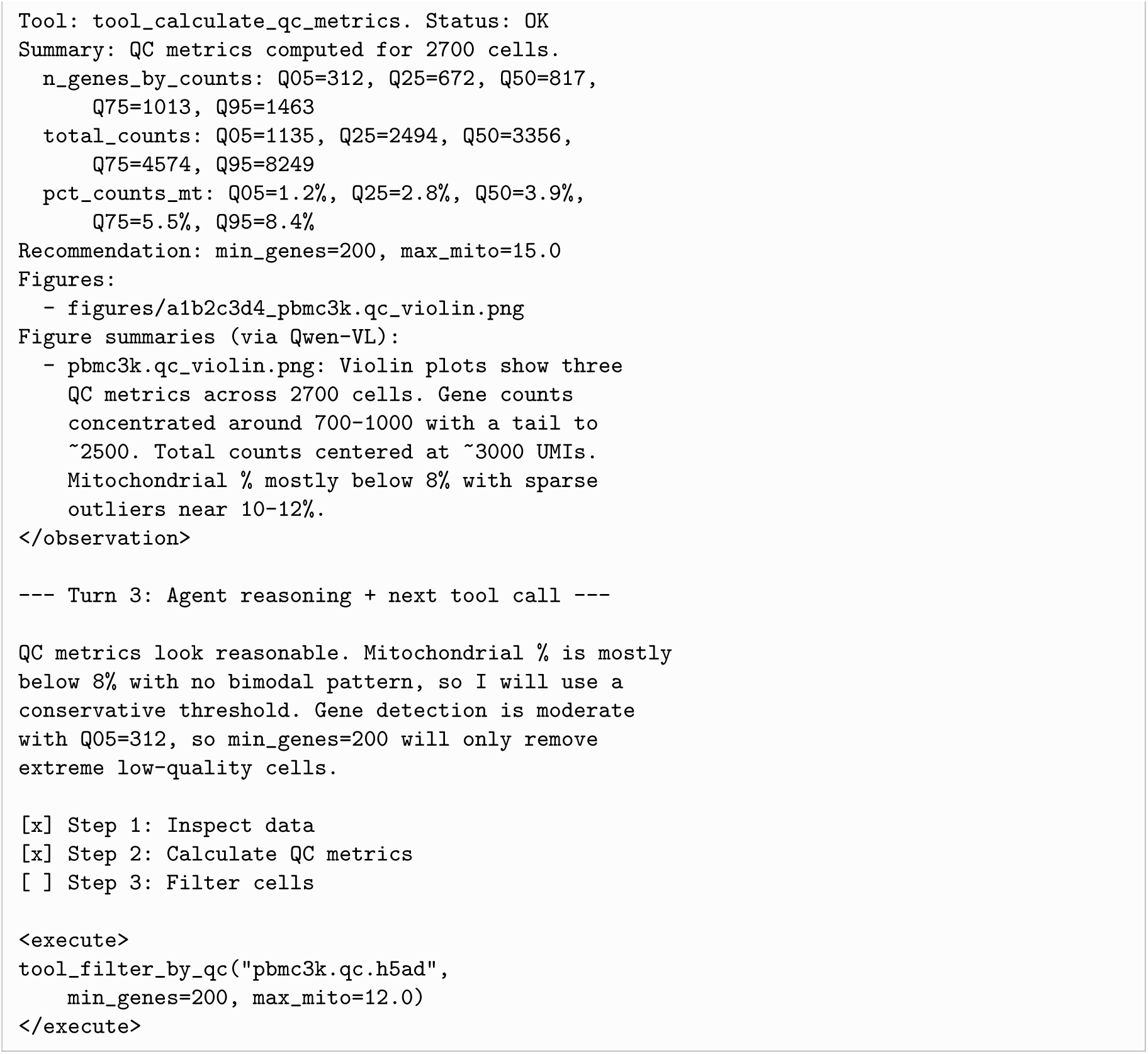

### 3.4 Self-Critic Prompt

After the agent emits a `<solution>` tag, the self-critic mechanism, when enabled, injects a review prompt into the conversation and invokes the language model for one additional reasoning pass. The review prompt is appended as a user message following the pruned conversation history, with the original system prompt retained. The verbatim self-critic prompt is shown in Box 3.

The critic operates under the same XML protocol as the main agent: it may emit a new `<execute>` tag to trigger corrective tool calls, or confirm completion with a `<solution>` tag.

#### Box 3.

**Self-critic review prompt (verbatim)**

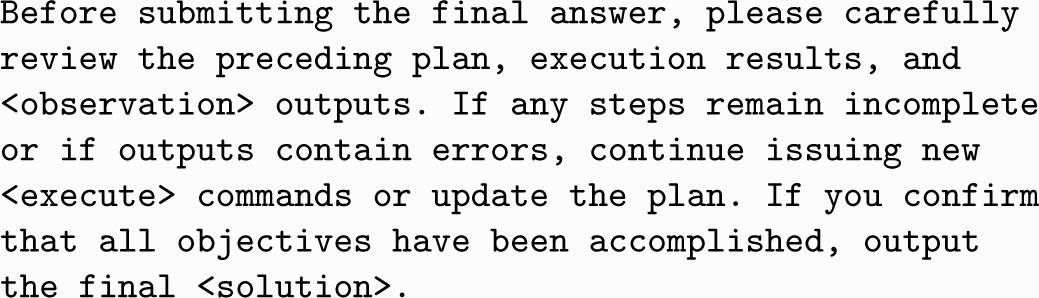

### 3.5 Vision-Language Model Prompt

When a tool generates a visualization, the VLM module (Supplementary Note 4) receives the image along with a fixed instruction prompt designed to elicit quantitative, actionable observations rather than generic descriptions. The verbatim VLM prompt is shown in Box 4.

The prompt explicitly requests measurable details because the main agent, which processes VLM summaries as text, benefits more from numeric observations that can guide parameter choices than from purely qualitative descriptions. The 100-word limit is enforced both through the prompt instruction and through post hoc truncation in the agent code.

#### Box 4.

**VLM image summary prompt (verbatim)**

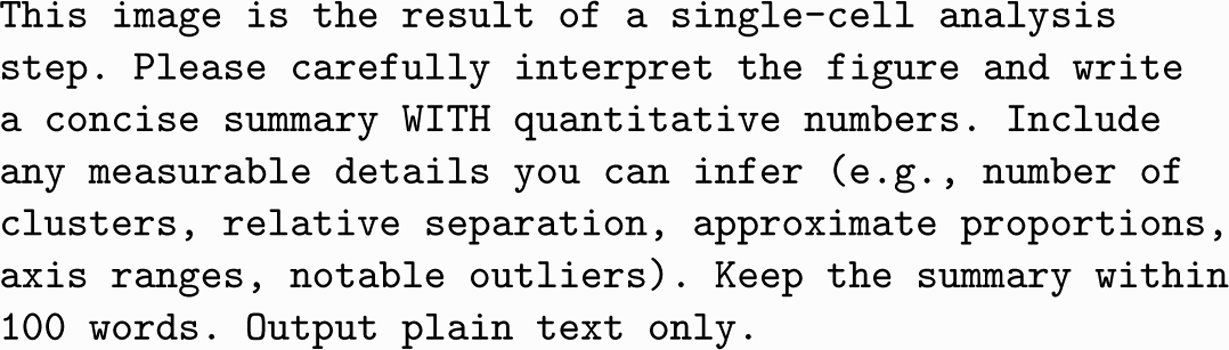

### 3.6 Prompt Design Rationale

Several design choices in the prompt system merit explicit justification, as they reflect trade-offs specific to piloting compact language models through multi-step analytical workflows.

#### Redundant constraint encoding

The preprocessing order (normalize `→` log `→` HVG `→` scale `→` PCA) is stated repeatedly in the system prompt, including a numbered sequence, a negative rule, and a worked code example. This redundancy is intended to reduce order violations in long multi-step sessions.

#### Checklist-driven planning

The prompt requires the agent to produce an explicit analysis plan with checkboxes in its first response and to mark each item as completed or skipped with justification before emitting a `<solution>`. This externalizes progress tracking into the conversation history.

#### Data-conditional threshold guidance

Rather than prescribing fixed QC thresholds, the prompt provides a mapping from observed quantile statistics to plausible threshold ranges. This allows the agent to adapt parameter choices to the structure of the input dataset.

#### Single-execute constraint

Restricting the model to one `<execute>` block per turn forces the agent to observe each tool output before planning the next step, producing a cleaner thought-action-observation loop.

### Supplementary Note 4: Vision-Language Model Integration

This note documents the design and operation of the vision-language model (VLM) module that enables CellPilot to incorporate visual evidence into its reasoning process. Section 4.4 of the main text introduces the rationale for multimodal observation enrichment and identifies the VLM backbone model. Here we describe the technical integration, output handling, and provide a concrete example from a benchmark annotation session.

#### 4.1 Module Architecture

The VLM module is implemented as a separate `QwenVisionSummarizer` class that is instantiated lazily upon first use rather than at agent initialization. When a tool execution produces one or more image files, the `execute` node in the agent graph passes the image paths to the summarization method, which coordinates VLM inference. Each image is loaded from disk, base64-encoded, and sent to the VLM alongside a fixed instruction prompt via an OpenAI-compatible API endpoint. The returned text summary is appended directly to the `<observation>` block as a `Figure summaries` section, making visual evidence available to the language model in the same context as the numeric tool output. Supported input formats include PNG, JPEG, and WebP.

#### 4.2 Output Constraints and Computational Cost Control

Two mechanisms limit the computational cost and context consumption of VLM summaries. First, each tool invocation triggers summarization of at most 2 images, even when more output figures are generated. Images are processed in the order they appear in the tool output. Second, each summary is constrained to 100 words through both the prompt instruction and post hoc truncation enforced in the agent code. These constraints allow VLM summaries to enrich the observation context without dominating the available context window.

#### 4.3 Caching and Failure Handling

VLM summarization operates on a best-effort basis. Summaries are cached per file using a composite key derived from the absolute file path, last modification timestamp, and file size, preventing redundant VLM calls when the same visualization is referenced repeatedly. If VLM inference fails for any reason, an empty result is returned silently rather than injecting error text into the `<observation>` block. Failed summaries are logged to the session directory for diagnosis but are not exposed to the main agent. VLM invocation is gated on the presence of a valid API key; when no key is configured, the module is disabled and the agent proceeds with numeric observations only.

#### 4.4 Example VLM-Guided Decision Making

To illustrate the VLM module in practice, we present an example from a CellPilot annotation session on a Tabula Sapiens kidney dataset. After differential expression analysis and marker gene cross-referencing, `tool_query_markers` produced a marker hit heatmap visualizing the number of DEG-marker overlaps between each cluster and each candidate cell type. Box 1 shows the heatmap (right) alongside the verbatim VLM summary (left) as it was appended to the agent’s `<observation>` context.

##### Box 1.

**VLM input and output: marker hit heatmap (Kidney, Tabula Sapiens)**

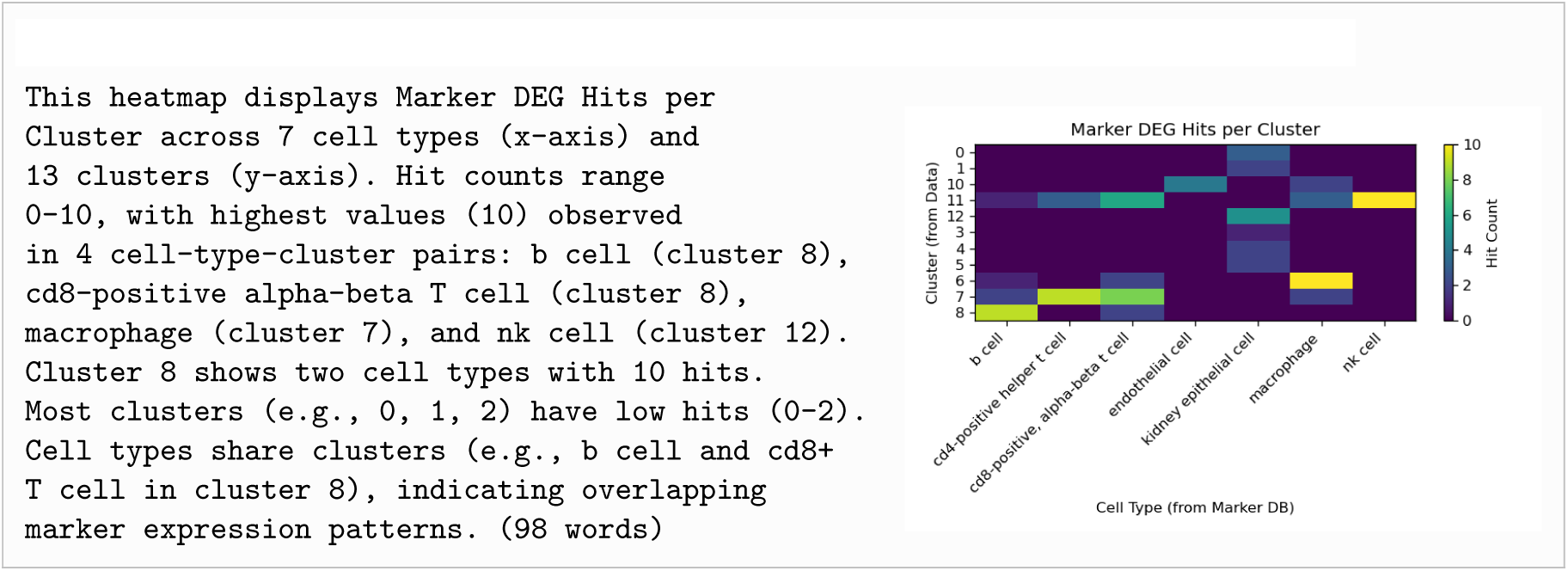

This summary provided the agent with three actionable observations: a subset of clusters showed dominant marker support for a single cell type, at least one cluster displayed ambiguous support across closely related lymphocyte labels, and several clusters showed low hit counts overall, suggesting limited marker database coverage for the dominant epithelial populations. This kind of compressed interpretation can help the agent decide whether additional refinement steps are warranted.

### Supplementary Note 5: Benchmark Design, Dataset Preparation, and Fairness Controls

Section 4.6 of the main text describes the benchmark datasets and the general principle that unified inputs were used across methods without dataset-specific tuning. This note provides the technical details of dataset organization, per-method input assumptions, cluster-level evaluation alignment, and fairness controls that are necessary for interpreting the comparative benchmark.

#### 5.1 Benchmark Inputs and Per-Tissue Evaluation Units

Each benchmark dataset was partitioned into per-tissue subsets, and each tissue was treated as an independent evaluation unit. For CellPilot, the input to each session consisted of a raw count matrix for a single tissue together with a marker gene list or differential expression gene list when available. Starting from this input, CellPilot autonomously executed quality control, normalization, highly variable gene selection, principal component analysis, neighbor graph construction, Leiden clustering, differential expression analysis, and annotation without manual intervention.

This per-tissue design reflects a practical single-cell analysis setting in which one tissue is analyzed at a time, and it also allows tissue-level scores to be computed independently.

#### 5.2 Shared Dataset Subsets Across Methods

All methods were evaluated on the same per-tissue subsets drawn from the same benchmark datasets. No method received a privileged subset of tissues or cells. Likewise, no dataset-specific prompt editing or manual correction was introduced for CellPilot. The same global prompt template and the same tool set were used throughout the benchmark. Parameter values used by CellPilot were selected by the agent itself on the basis of observed data characteristics within each session rather than by human tuning.

#### 5.3 Benchmark Datasets and Format Standardization

Expression matrices for all three datasets were downloaded from their original public repositories. GTEx single-cell and single-nucleus data were obtained from the GTEx Portal (https://gtexportal.org; PMID: 35549429). Tabula Sapiens data were downloaded from the CZI cellxgene portal (https://cellxgene.cziscience.com; PMID: 35549404). Mouse Cell Atlas data were retrieved from the MCA website (http://bis.zju.edu.cn/MCA/; PMID: 29474909; GEO accession: GSE108097).

All datasets were converted to AnnData (.h5ad) format prior to analysis. For GTEx and Tabula Sapiens, where data were already distributed as AnnData objects, conversion consisted of extracting the raw count layer and discarding precomputed embeddings or transformed representations so that CellPilot operated on count-level inputs. For MCA, the original Microwell-seq count matrices were loaded into AnnData structures with gene names as variable indices and cell barcodes as observation indices. No gene filtering, cell filtering, or normalization was applied before CellPilot processing.

#### 5.4 Tissue-Level Organization

Each dataset was split into per-tissue subsets, with each tissue processed as an independent CellPilot session. Supplementary Table 2 summarizes the key characteristics of each dataset, including species, tissue count, cell type count, and publication references.

GTEx comprises 7 tissues (Breast, Esophagus, Heart, Lung, Prostate, Skeletal Muscle, Skin), with cell counts per tissue ranging from approximately 10,000 to 50,000 and 41 distinct cell types across all tissues.

Tabula Sapiens comprises 24 tissues spanning immune (Blood, Bone Marrow, Lymph Node, Spleen, Thymus), epithelial (Lung, Large Intestine, Small Intestine, Bladder, Trachea), solid (Heart, Liver, Kidney, Pancreas), glandular (Mammary, Prostate, Salivary Gland), and other organ systems (Eye, Fat, Muscle, Skin, Tongue, Uterus, Vasculature), encompassing 171 distinct cell types.

MCA spans 51 tissues with 64 cell types. Unlike GTEx and Tabula Sapiens, MCA tissues vary considerably in sequencing depth and cell recovery, with some tissues containing fewer than 500 cells. These small-tissue samples present particular challenges for clustering and differential expression analysis.

#### 5.5 Baseline-Specific Input Assumptions

The four baseline methods evaluated in the main benchmark differ substantially in the upstream information they receive as input. This distinction is important for interpreting comparative performance, as methods that are provided with ground-truth cluster assignments effectively operate under an oracle clustering assumption, removing a major source of analytical uncertainty. CellPilot, by contrast, receives only the raw count matrix and determines all upstream analytical decisions autonomously, without access to ground-truth cluster boundaries or precomputed marker gene lists (Methods). A summary of the input requirements for each method is provided in Supplementary Table 1.

Among the baselines, SingleR operates without access to ground-truth cluster boundaries:

##### SingleR

SingleR operates at the single-cell level, assigning labels by comparing each cell independently to a reference expression atlas using correlation-based similarity. Cluster assignments are not required for prediction. In this study, predictions were aggregated to the cluster level by majority vote for evaluation, but all labels were generated independently at the single-cell level.

The remaining three baselines require ground-truth cluster assignments as input:

##### ScType

ScType operates at the cluster level and does not perform clustering. It takes predefined cluster assignments together with expression data and assigns a label to each cluster using the ScType marker database. In this benchmark, the supplied cluster assignments correspond to the ground-truth labels provided with each dataset.

##### GPTCelltype4

GPTCelltype4 also operates at the cluster level. It takes marker gene lists derived from ground-truth clusters, together with tissue information, and returns predicted cell type labels using GPT-4. Both the cluster definitions and the corresponding differential expression results were derived from the ground-truth annotations in the benchmark datasets.

##### CASSIA

CASSIA operates at the cluster level using a multi-step prompting strategy with GPT-4o. Similar to GPTCelltype4, it relies on marker gene lists derived from ground-truth clusters, along with tissue context information. Default parameters were used throughout, consistent with the other baseline methods.

#### 5.6 Shared Inputs and Clustering Assumptions Across Methods

To clarify the use of “same preprocessed clusters” in the main text, it is important to distinguish between their role as method inputs and as the common reference for evaluation.

For clustering, ScType, GPTCelltype4, and CASSIA were provided with the ground-truth cluster assignments from the benchmark datasets as direct input. In contrast, SingleR does not use cluster assignments for prediction, and CellPilot performs clustering from raw data. Neither SingleR nor CellPilot had access to the benchmark-provided cluster boundaries during annotation. For evaluation, however, predictions from all methods were mapped to the same ground-truth clusters to ensure a consistent basis for comparison.

For marker gene inputs, GPTCelltype4, CASSIA, ScType, and CellPilot all used the top 10 differentially expressed genes per cluster as the basis for annotation. For the three cluster-dependent baselines, these marker gene lists were precomputed from the ground-truth clusters. CellPilot instead computed differential expression from its own clusters, while maintaining the same marker list size for consistency. SingleR does not rely on marker genes and instead performs direct expression profile matching against a reference atlas.

#### 5.7 Cluster-Level Evaluation of CellPilot

Because CellPilot performs clustering by itself, its inferred clusters do not necessarily align with the reference clusters provided in the benchmark datasets. To enable direct comparison with other methods, CellPilot predictions were therefore mapped to the dataset-defined reference clusters for evaluation.

Two complementary aggregation schemes were used.

##### CellPilot (Score)

Each cell was assigned an agreement score based on the comparison between its predicted label and the reference annotation. For each reference cluster, these cell-level scores were averaged to obtain a continuous cluster-level score ranging from 0 to 1.

##### CellPilot (Vote)

For each reference cluster, the most frequent predicted cell type among its constituent cells was taken as the cluster-level label. This label was then compared with the reference annotation using the three-level scoring scheme described in Supplementary Note 6, yielding a score of 0, 0.5, or 1.

At the tissue level, scores were computed as the unweighted mean across all reference clusters within each tissue, such that each cluster contributed equally regardless of its size.

#### 5.8 Cluster Aggregation Policy and Absence of Majority Thresholding

No additional majority threshold was imposed when summarizing predictions within reference clusters. In particular, clusters that contained heterogeneous cell populations were not excluded, relabeled, or separated on the basis of purity criteria. All reference clusters were retained in evaluation as provided by the benchmark datasets.

#### 5.9 Ground-Truth Label Harmonization

Predicted and ground-truth cell type labels were mapped to Cell Ontology names and broad cell type categories to enable standardized evaluation. For Tabula Sapiens, the finest available annotation level (`cell_ontology_class`) was used as the primary reference label. For MCA, the original study annotations were used. Cross-species evaluation in MCA was performed without explicit mouse-to-human ortholog conversion at the input stage, meaning that the method had to rely on conserved marker patterns rather than direct gene-name remapping.

#### 5.10 Marker Gene Lists for GPTCelltype4 Benchmarking

To ensure comparability with GPTCelltype4, marker gene lists were generated following the protocol described by Hou and Ji for the corresponding datasets. For GTEx, raw counts were library-size normalized and log-transformed after adding a pseudocount of 1, ComBat was applied to account for protocol- and sex-specific batch effects, and Welch’s `t`-test was used to identify differential genes. For Tabula Sapiens, raw counts were library-size normalized and log-transformed after adding a pseudocount of 1, and Seurat’s `FindAllMarkers()` with default settings was used to obtain differential genes within each tissue. For MCA, the differential gene lists provided by the original study were used. In the primary benchmark experiments, GPTCelltype4 was provided with the top 10 marker genes per cluster as recommended in the original study.

#### 5.11 No Manual Intervention

No dataset-specific prompt engineering, no tissue-specific parameter presets, and no manual post hoc correction of CellPilot predictions were applied during benchmarking. CellPilot used a unified prompt and made tool-parameter decisions on the basis of session-specific observations. This design was chosen to evaluate the method as a general-purpose autonomous framework rather than as a manually tuned workflow.

### Supplementary Note 6: Marker Database Construction, Annotation Provenance, and Hierarchical Evaluation

This note describes the origin of the marker information used by CellPilot, the scoring logic implemented in the marker-query step, the conditions under which clusters are designated as unassigned, and the hierarchical rules used to classify prediction outcomes as full matches, partial matches, or mismatches.

#### 6.1 Marker Source Hierarchy

CellPilot first allows the user to provide a custom marker gene list. This is the preferred mode when the user has access to a curated tissue-specific or study-specific marker resource. When no user-supplied marker file is available, the system falls back to an internal default marker database derived from PanglaoDB.

This design separates two use cases. In one setting, CellPilot serves as a flexible annotation framework that can incorporate expert-provided prior knowledge. In the other, it operates in a default mode using a broad public marker resource.

#### 6.2 Marker Query Procedure

The core marker-based annotation logic is implemented in the `tool_query_markers` function in `marker_db.py`. For each cluster, the tool extracts the top *K* differentially expressed genes from `rank_genes_groups`, with `top_k=50` by default. Gene symbols are converted to uppercase before matching to the marker database. If a queried gene is found in the marker table for one or more cell types, the hit count for the corresponding cell type is incremented. This procedure yields, for each cluster, a dictionary of the form `{cell_type: hit_count}`.

The predicted cell type for the cluster is defined as the cell type with the highest hit count.

#### 6.3 Marker-Based Confidence Score

The confidence score used in the primary marker-based annotation layer is defined as

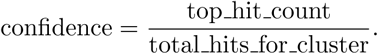

Thus, if a cluster produces 20 total marker hits across all candidate cell types and 15 of those hits support a T-cell label, the resulting confidence score is 15/20 = 0.75. This score reflects the concentration of marker evidence on the top-supported label rather than the absolute number of matched markers.

#### 6.4 Triggering of Unassigned Labels

A cluster is designated as `Unassigned` only when none of its top *K* differentially expressed genes match any entry in the marker database. In implementation terms, this occurs when the cluster has no entry in the top-hit summary map and is later filled with `Unassigned` and a confidence of 0.0. The trigger is therefore strict: zero marker hits. If even a single DEG matches the marker database, the cluster receives a predicted label.

#### 6.5 Annotation Provenance and LLM Refinement

Marker-based annotation constitutes the primary annotation layer in CellPilot. Clusters labeled `Unassigned` are then eligible for a secondary refinement step using `tool_llm_annotate_unassigned`. The refined label, confidence score, and brief reasoning returned by the language model are stored separately from the original marker-based fields. This preserves annotation provenance and makes it possible to distinguish direct marker-supported assignments from LLM-based fallback assignments in downstream analyses.

#### 6.6 Scope and Limitations of the Marker Database

The default PanglaoDB-derived marker database provides broad coverage across many canonical cell types, but it is not tissue-complete and does not guarantee uniform resolution across all lineages. In particular, rare epithelial subtypes, transitional stromal populations, and species-specific cellular states may be underrepresented. For this reason, low marker hit counts or `Unassigned` outcomes do not necessarily indicate that a cluster lacks biological identity. Instead, they may reflect incomplete marker coverage in the reference database.

#### 6.7 Hierarchical Evaluation Rules

Each identified cell type annotation was assigned an unambiguous cell ontology name and, when applicable, a broad cell type name. A pair of identified cell type annotations was classified as *fully matched* if they shared the same annotation term or the same available Cell Ontology name. A pair was classified as *partially matched* if they had the same broad cell type name, or if one was subordinate to the other at the broad cell type level, but the detailed annotation term and Cell Ontology name differed. A pair was classified as a *mismatch* if broad cell type name, annotation term, and Cell Ontology name were all different.

This three-level scheme was used throughout the main benchmark to translate biologically near-miss predictions into an intermediate score rather than treating all non-identical labels as equally incorrect.

#### 6.8 Parent–Child Adjudication Examples

The partial-match category is intended to capture biologically related labels that differ in granularity. For example, a prediction of *stromal cell* for a reference label *fibroblast*, or the reverse, is treated as a partial match rather than a complete mismatch. Likewise, predictions at a broader lineage level can receive intermediate credit when they remain within the same biological compartment as the reference annotation.

#### 6.9 Broad Cell Type Mapping

Broad cell type categories were used both in hierarchical scoring and in downstream stratified analyses. This layer allows performance to be summarized at a coarser biological resolution, which is particularly helpful when comparing methods that differ in how often they assign broad parent labels versus fine-grained subtypes.

#### 6.10 Semantic Similarity Analysis with SapBERT

Semantic similarity analysis was used to further characterize residual errors beyond the categorical full/partial/mismatch scoring scheme. In particular, cases scored as partial matches or mismatches can still differ substantially in biological plausibility. SapBERT-based text embedding similarity therefore provides an auxiliary view of whether predicted and reference labels remain semantically close even when they are not identical under the hierarchical evaluation rules.

In the final manuscript version, this subsection will additionally document the exact input label normalization procedure, the pairwise comparison set used for the analysis, and the rationale for focusing on non-full-match cases in the semantic similarity plots.

### Supplementary Note 7: Cell-level Annotation Performance Across Annotation Granularity, Broad Cell Types, Cluster Size, and Tissue Context

#### 7.1 Annotation Granularity

Annotation performance differed modestly but consistently between major cell type and subtype annotations. Ground-truth labels classified as major cell types achieved a full-match rate of 83%, while subtype-level labels reached 78%. This difference is not unexpected, because many cell subtypes share a large proportion of their expressed gene repertoire with closely related populations. Naive and central memory T cells, for example, can be distinguished only by a relatively small number of differentially expressed genes, many of which are co-expressed at low levels across both populations. The retention of a 78% full-match rate at the subtype level suggests that CellPilot preserves annotation specificity at finer granularity more effectively than methods that systematically collapse uncertain cases to broad parent categories.

#### 7.2 Broad Cell Type Performance and Misclassification Patterns

To characterize annotation performance across major biological lineages independently of the original annotation granularity, all ground-truth labels were mapped to their corresponding broad cell type categories. The ten most frequently represented broad types in human datasets collectively covered 92.3% (581,535 of 629,934) of annotated cells; in mouse datasets, the equivalent coverage was 77.1% (45,687 of 59,250 cells) (Supplementary Fig. 4).

Among human lineages, cell types with highly specific and well-validated marker gene signatures performed best. Fibroblast (*n* = 56,781), Endothelial cell (*n* = 51,430), and B cell (*n* = 51,924) all showed consistently high full-match rates, consistent with the structural distinctiveness of their transcriptional profiles. Stem cell presented a more complex picture: 18% of stem cells were annotated as Epithelial cell, likely reflecting the epithelial-like gene expression programs retained by many tissue-resident progenitor populations. Granulocyte showed 14% misclassification to Macrophage/-monocyte, a pattern attributable to overlap in inflammatory gene modules shared across myeloid lineages (Supplementary Fig. 5a).

In mouse datasets, Stem cell (*n* = 4,428) and Stromal cell (*n* = 7,515) were annotated with high accuracy. Myeloid populations were more problematic: 21% of Macrophage/monocyte cells were assigned to Stem cell, a pattern that may reflect reduced conservation of canonical myeloid marker genes in the mouse reference and the resulting difficulty of distinguishing these populations in a cross-species annotation setting. Erythroid cell showed 18% misclassification to Muscle cell. Although the transcriptional basis of this pattern is less obvious, both lineages can share cytoskeletal and membrane-associated signatures, and the relatively small erythroid cluster sizes in some mouse tissues may further increase instability (Supplementary Fig. 5b).

Across both species, misclassification events were concentrated within biologically related lineages and rarely spanned unrelated identities, consistent with the semantic similarity patterns reported in the main text.

#### 7.3 Cluster Size Effects

The dependence of annotation accuracy on cluster size reflects a fundamental constraint of differential expression analysis. In clusters of ten or fewer cells, the statistical power to identify stable and reproducible marker rankings is limited, and the resulting DEG lists are more sensitive to sampling noise. Under these conditions, the genes passed to the annotation step may vary substantially across runs and may not faithfully represent the canonical marker structure of the underlying cell type. This instability propagates directly into annotation quality.

In practical terms, very small clusters warrant closer manual review regardless of the annotation tool used. CellPilot confidence scores offer one practical way to flag such cases for further inspection without requiring a fixed size cutoff in advance.

#### 7.4 Tissue-Level Performance

Agreement proportions varied considerably across tissue types in both Tabula Sapiens and HCA datasets (Fig. 4d). Tissues with predominantly haematopoietic compositions achieved the highest full-match rates: Tongue, Bone Marrow, Blood, and Lymph Node all showed strong annotation accuracy, consistent with the well-characterized and highly specific marker gene signatures of immune lineages. Esophagus in the HCA dataset performed similarly well.

Tissues with predominantly epithelial or stromal compositions showed higher proportions of partial matches. Mammary and Kidney were notable in this regard, reflecting the transcriptional similarity among epithelial and stromal subtypes that complicates boundary assignment at fine resolution. Skeletal muscle, Muscle, and Heart showed intermediate performance; muscle lineage markers are moderately specific, but subtype boundaries vary across tissue contexts and annotation schemes, contributing more often to partial rather than full agreement.

These tissue-dependent patterns support the practical use of confidence-based filtering rather than a single global threshold. Tissues that are intrinsically more difficult to annotate can be expected to produce lower-confidence predictions on average, allowing review effort to be allocated more selectively.

## Supplementary Tables

**Table 1.**
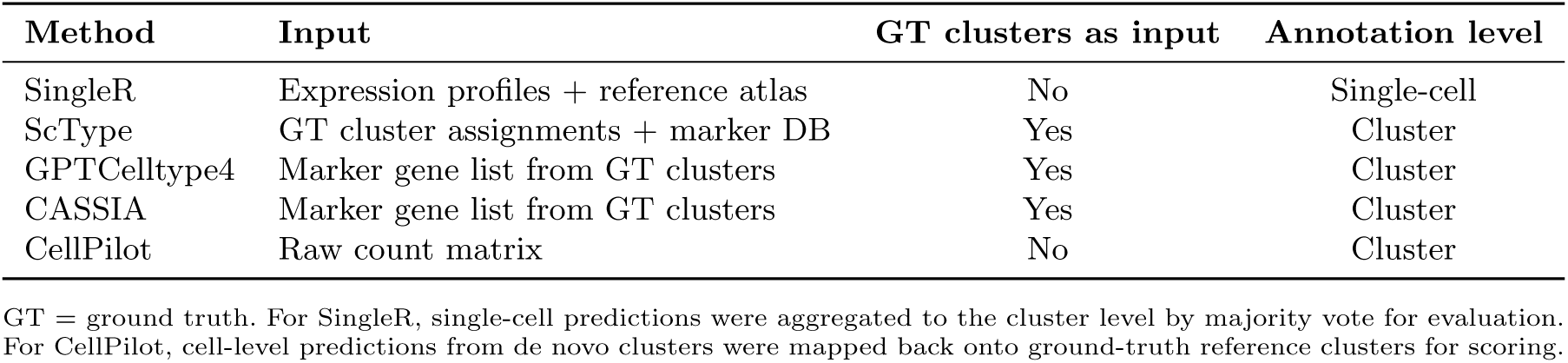
Summary of input requirements and clustering assumptions across annotation methods. Ground-truth cluster access indicates whether the method receives benchmark-provided cluster assignments as direct input for annotation. Evaluation unit for all methods is the ground-truth reference cluster.

## 1 Supplementary Figures

**Supplementary Fig. 1.**
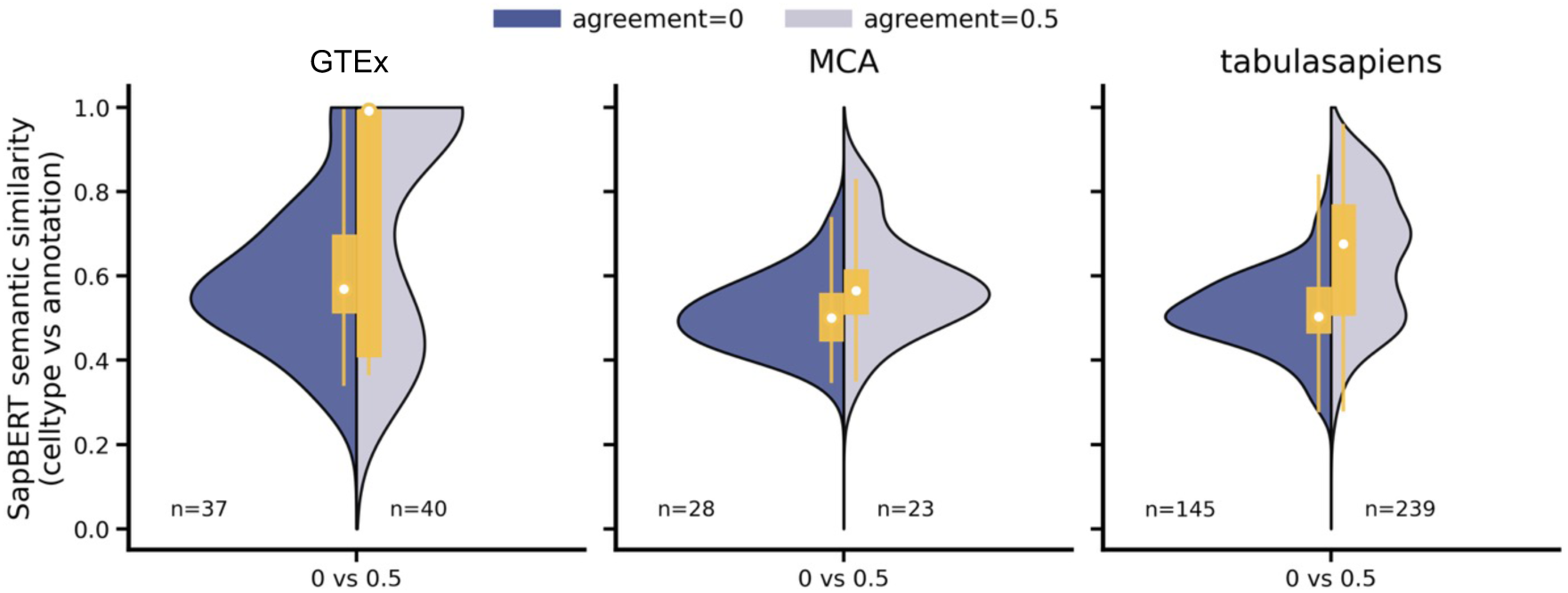
SapBERT-based semantic similarity between SingleR-predicted and true cell-type names for incorrect cases across GTEx, MCA, and Tabula Sapiens. Split violin plots compare completely incorrect predictions (agreement = 0) and partially correct predictions (agreement = 0.5). Violin width reflects score density, and inner boxes indicate the interquartile range with median. Higher values denote greater semantic proximity between predicted and true labels.

**Supplementary Fig. 2.**
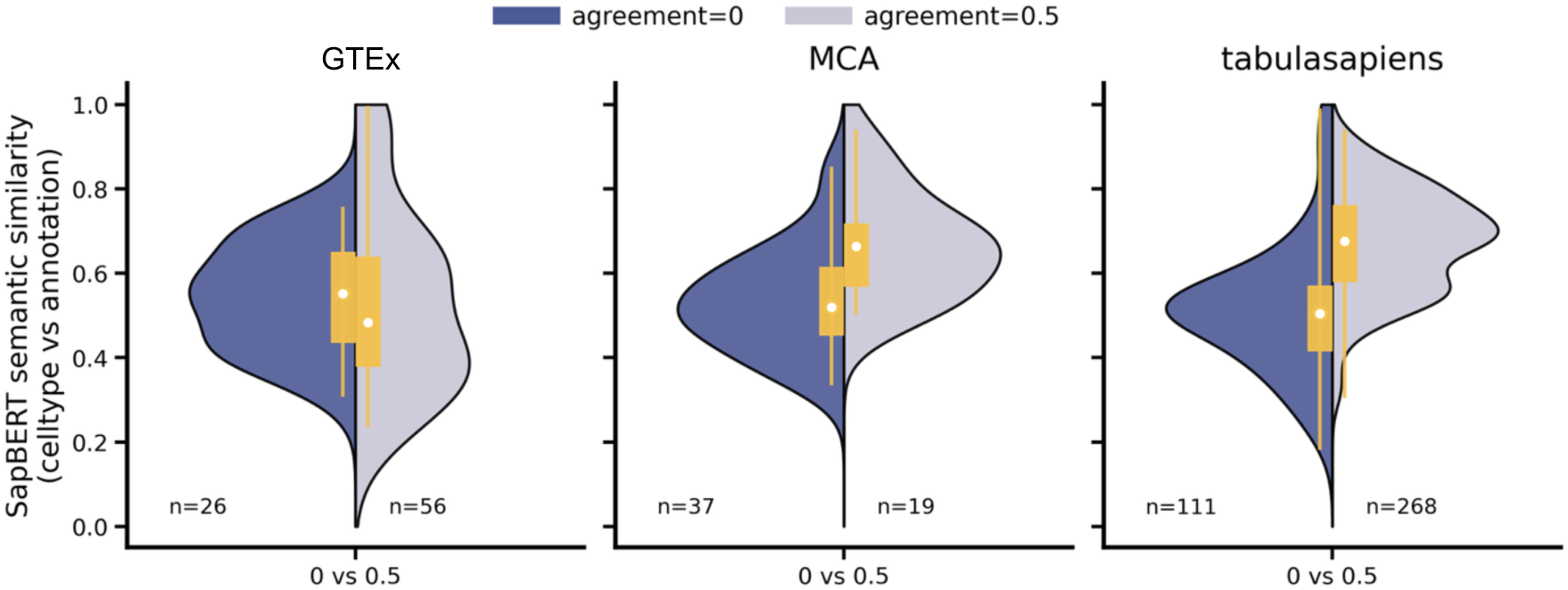
SapBERT-based semantic similarity between SCType-predicted and true cell-type names for incorrect cases across GTEx, MCA, and Tabula Sapiens. Split violin plots compare completely incorrect predictions (agreement = 0) and partially correct predictions (agreement = 0.5). Violin width reflects score density, and inner boxes indicate the interquartile range with median. Higher values denote greater semantic proximity between predicted and true labels.

**Supplementary Fig. 3.**
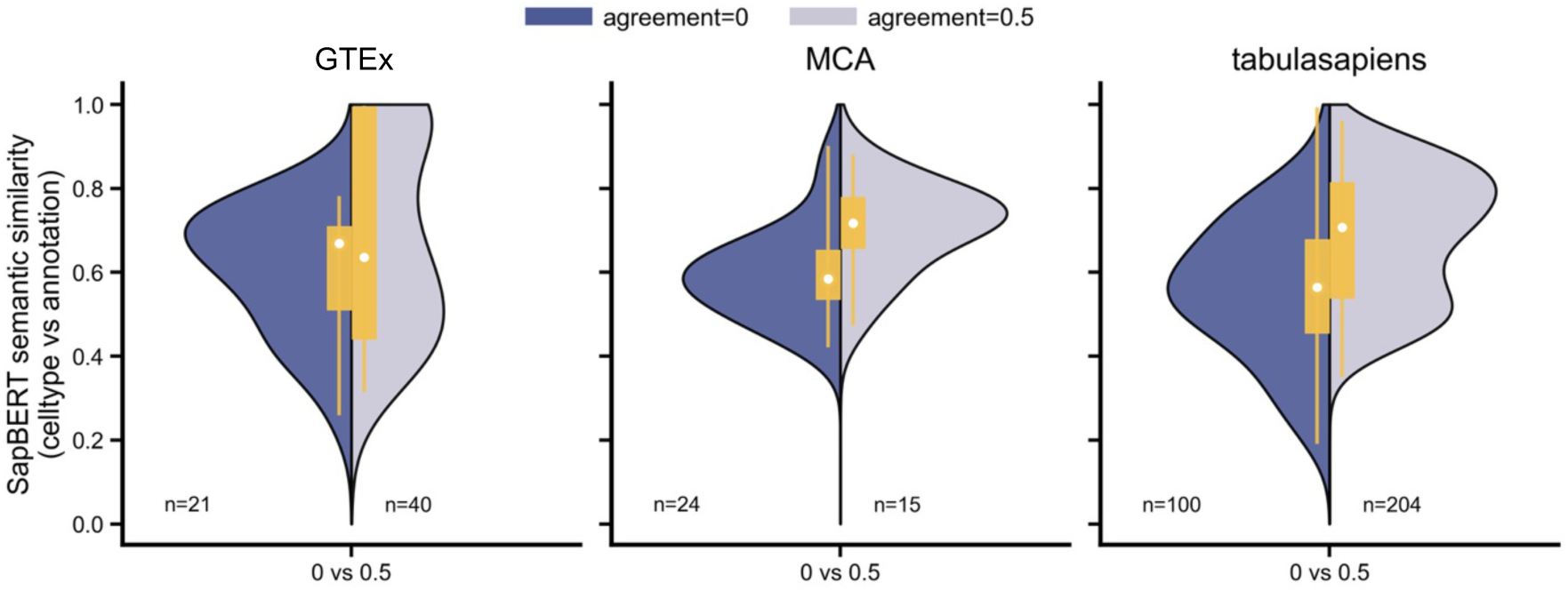
SapBERT-based semantic similarity between GPTCelltype4-predicted and true cell-type names for incorrect cases across GTEx, MCA, and Tabula Sapiens. Split violin plots compare completely incorrect predictions (agreement = 0) and partially correct predictions (agreement = 0.5). Violin width reflects score density, and inner boxes indicate the interquartile range with median. Higher values denote greater semantic proximity between predicted and true labels.

**Supplementary Fig. 4.**
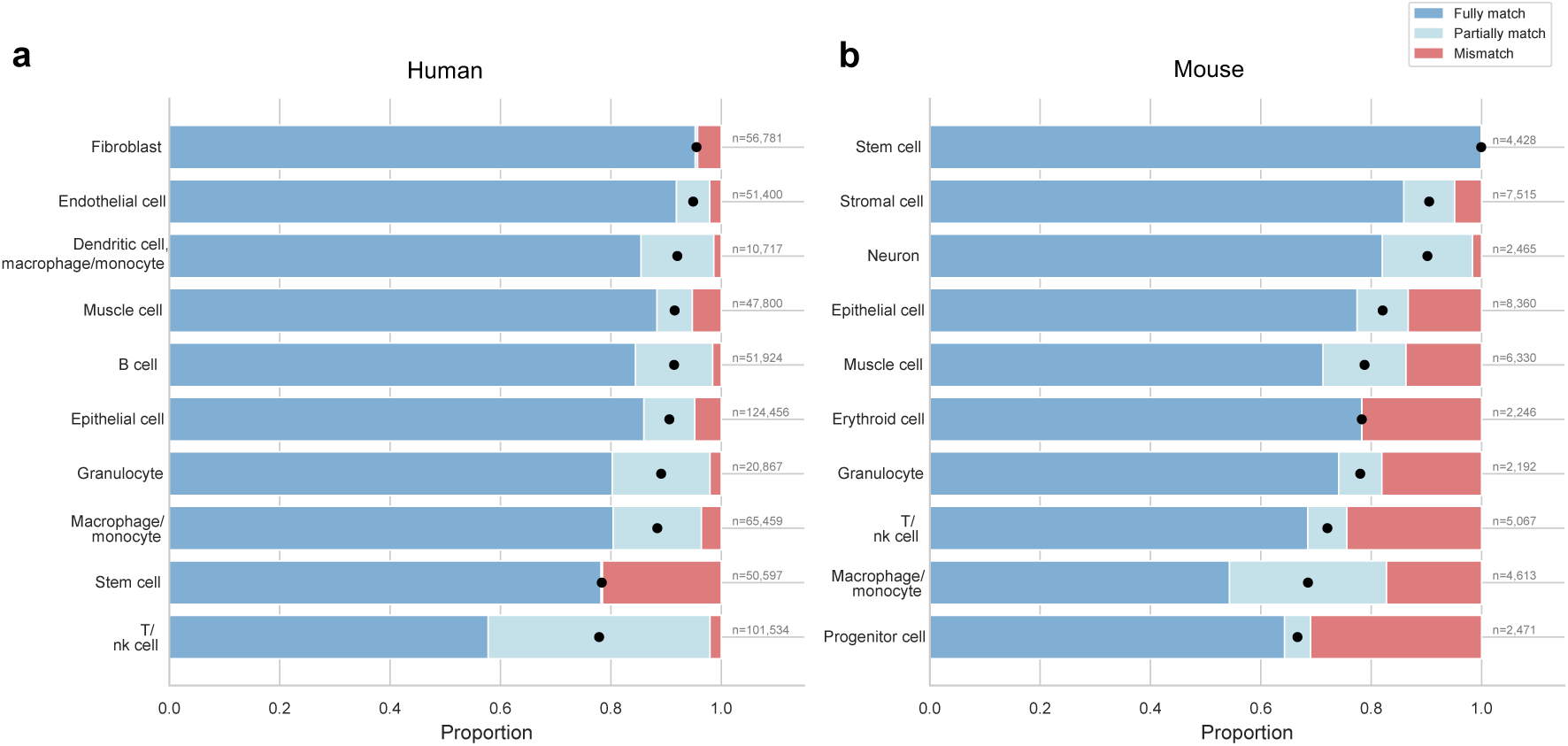
Top 10 broad cell types in human and mouse datasets. (a) Human datasets (GTEx + TS): For the ten most frequent broad cell types, the proportions of agreement scores (0, 0.5, and 1) are shown, representing completely incorrect, partially correct, and fully correct annotations, respectively. (b) Mouse datasets (MCA): The same analysis is presented for the top ten broad cell types in mouse. Numbers displayed on the right indicate the total number of cells (n) in each broad cell-type category.

**Supplementary Fig. 5.**
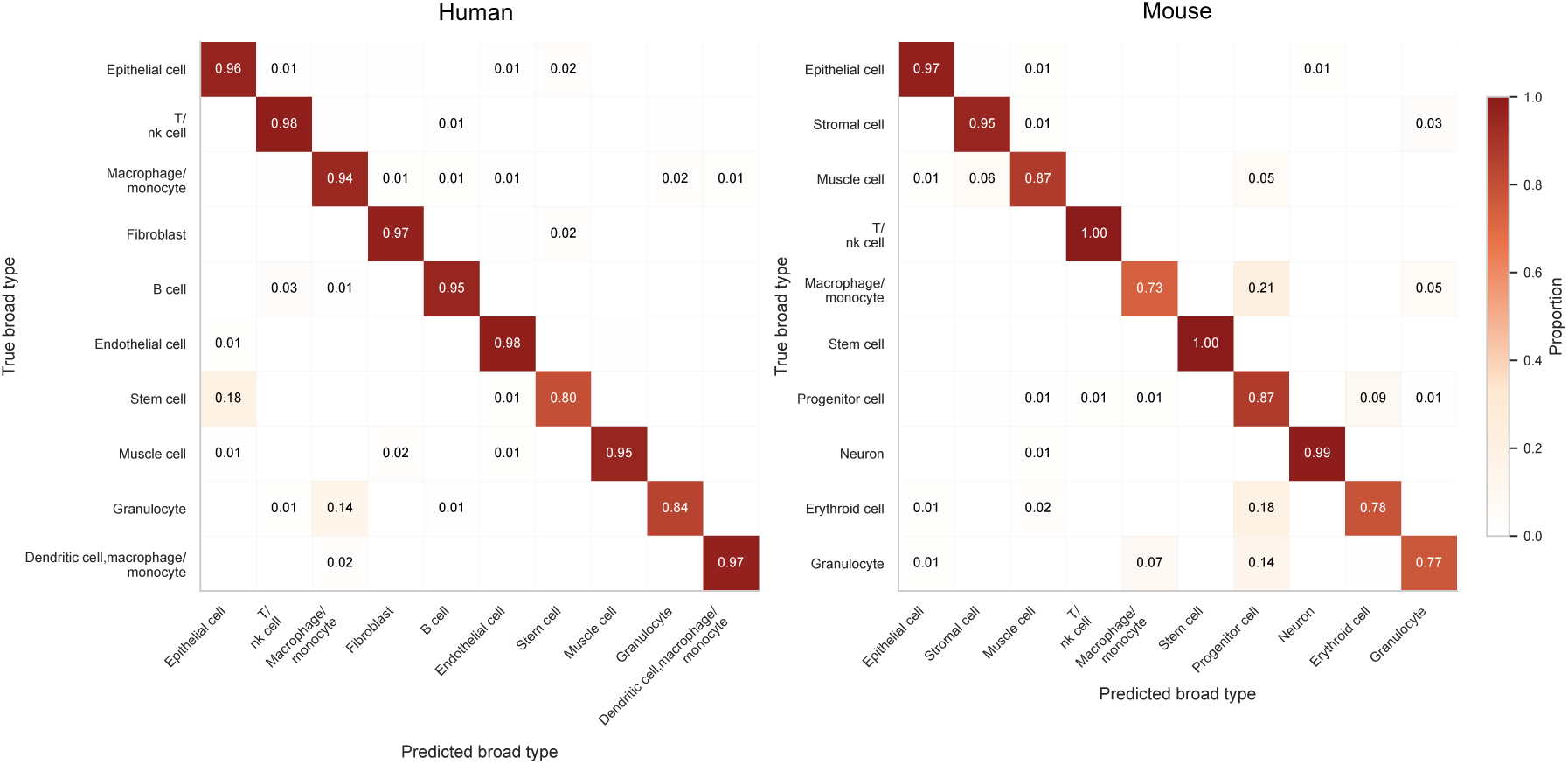
Broad-type confusion matrices for the top 10 cell categories in human and mouse datasets. (a) Human (GTEx + TS): Confusion matrix for the ten most frequent broad cell types. Rows indicate true broad cell types and columns indicate predicted broad cell types. Values represent the proportion of cells within each true category assigned to each predicted category. (b) Mouse (MCA): Confusion matrix for the top ten broad cell types in mouse, displayed in the same format. Diagonal entries correspond to correct broad-type assignments, while off-diagonal entries indicate misclassification patterns.

**Supplementary Fig. 6.**
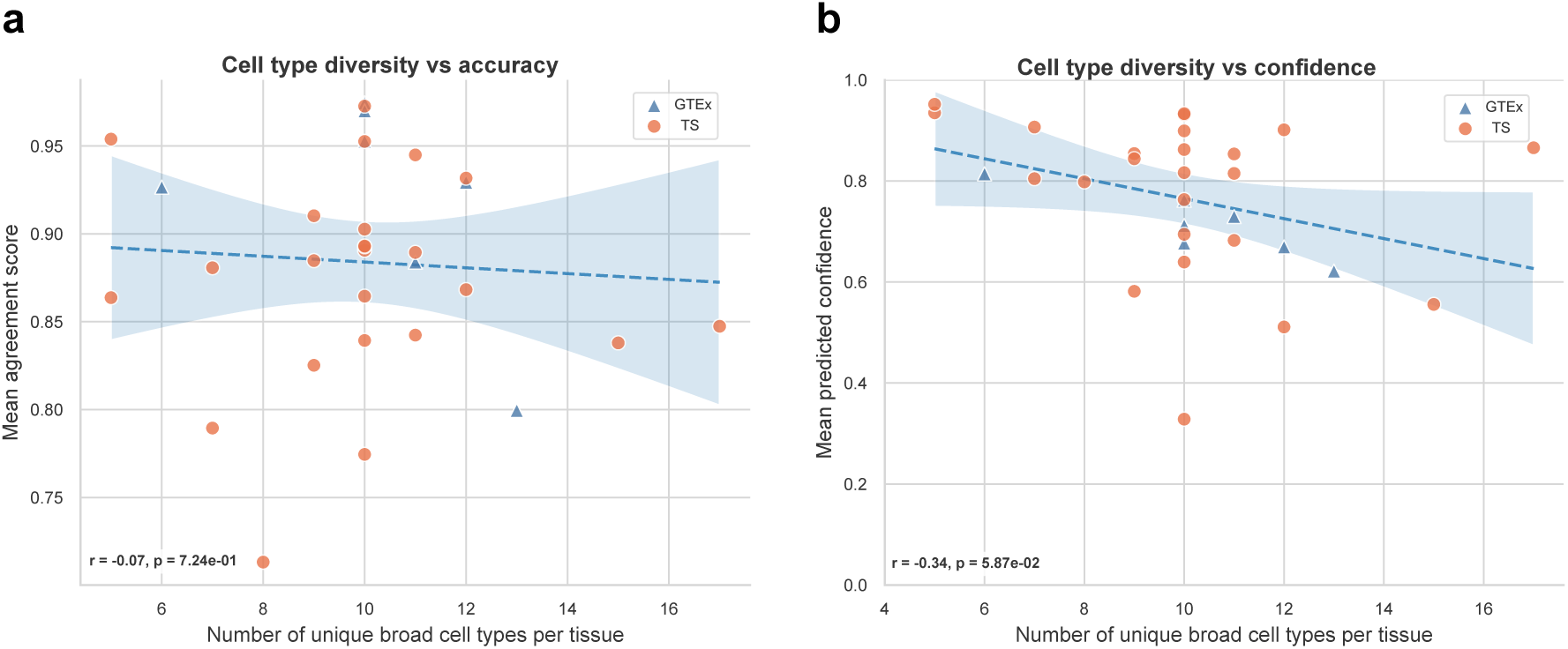
Relationship between cell-type diversity and model performance. (a) Association between the number of unique broad cell types per tissue and the mean agreement score. Each point represents a tissue from human datasets (GTEx, TS). The dashed line indicates the linear regression fit with shaded area representing the 95% confidence interval. The reported r and p values correspond to Pearson correlation. (b) Association between the number of unique broad cell types per tissue and the mean predicted confidence. Points and regression lines are shown as in (a). Negative trends suggest reduced confidence and slightly lower agreement in tissues with greater cell-type diversity.

**Supplementary Fig. 7.**
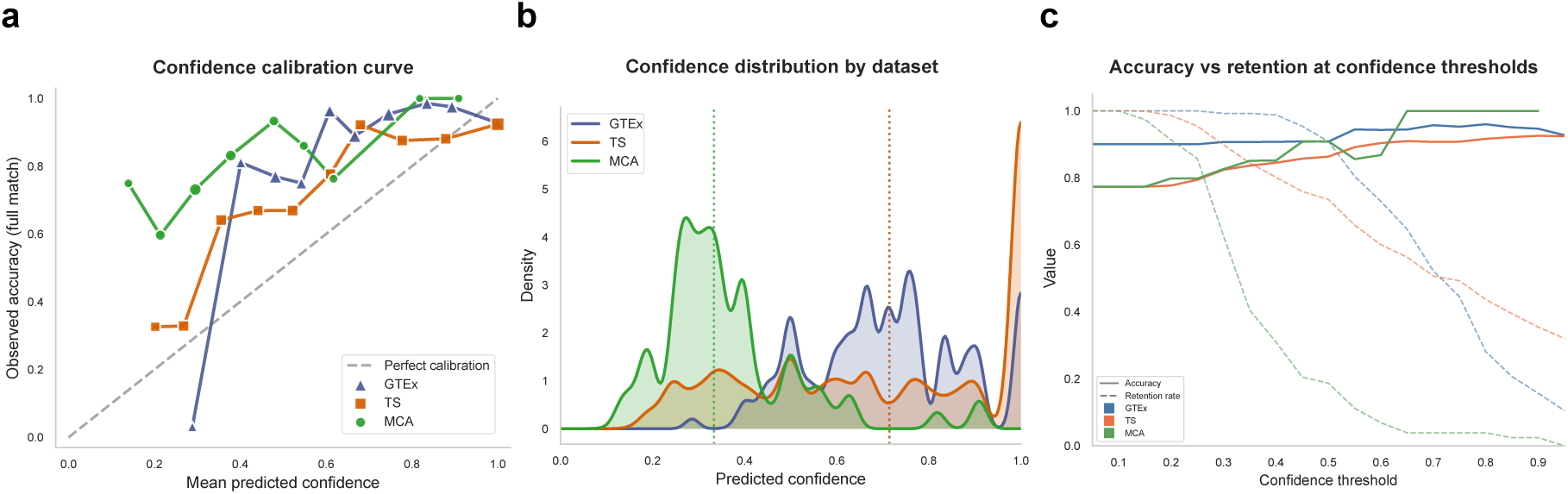
Evaluation of confidence calibration and reliability across datasets. (a) Confidence calibration curves for GTEx, TS, and MCA. The x-axis represents the mean predicted confidence within bins, and the y-axis shows the observed full-match accuracy. The dashed diagonal line indicates perfect calibration. Points above the diagonal reflect underconfident predictions, whereas points below indicate overconfidence. (b) Distribution of predicted confidence scores for each dataset. Curves represent kernel density estimates of confidence values. Vertical dotted lines indicate the median confidence per dataset. (c) Accuracy–retention trade-off across confidence thresholds. Solid lines show observed accuracy among predictions with confidence greater than or equal to each threshold, while dashed lines represent the corresponding retention rate (fraction of predictions retained). This panel illustrates how increasing the confidence threshold improves accuracy at the cost of coverage.

**Supplementary Fig. 8.**
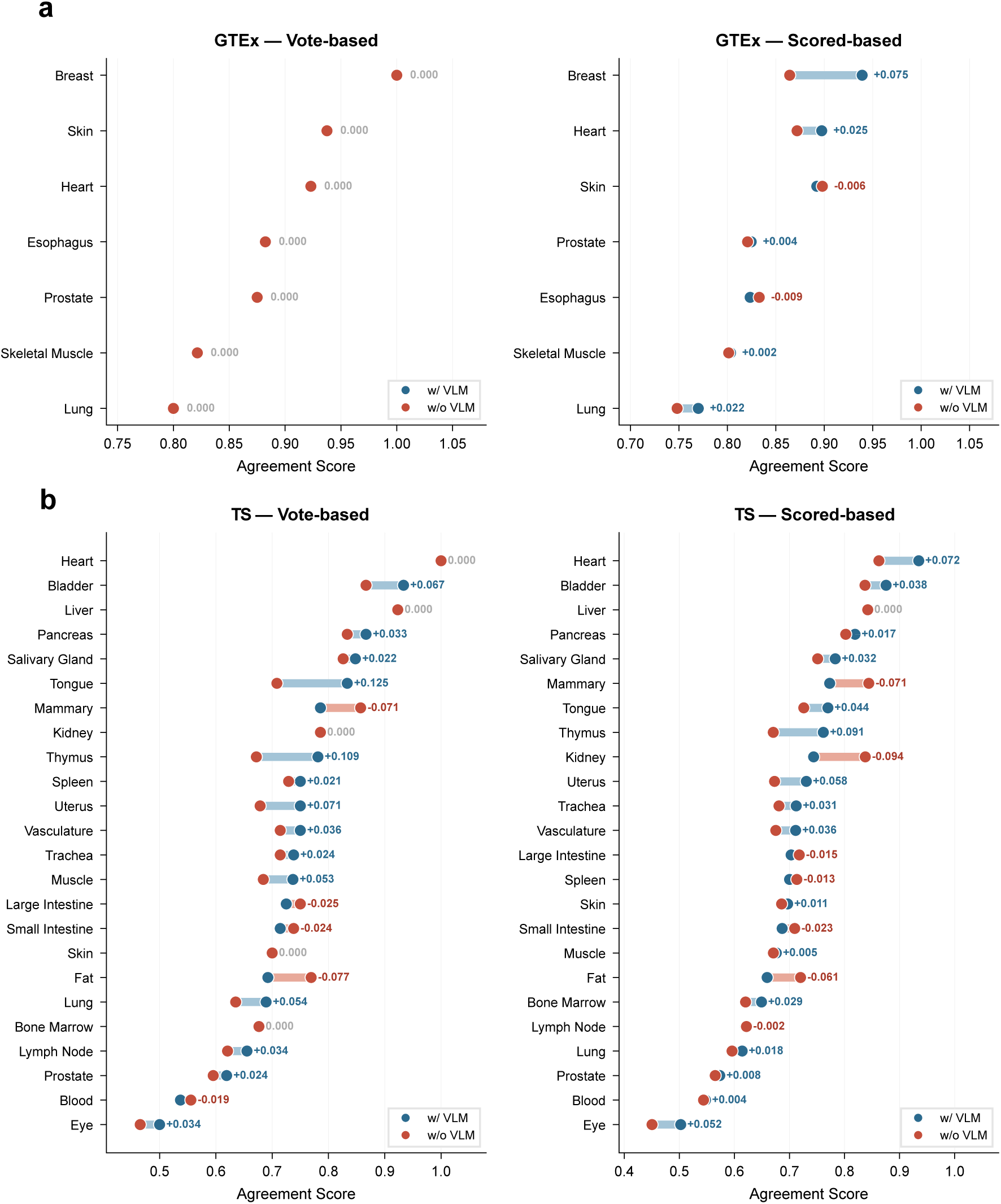
Impact of vision-language model (VLM) integration on tissue-level annotation agreement. (a)Agreement scores with and without VLM across GTEx tissues using vote-based (left) and score-based (right) consensus strategies. Numerical annotations denote the score difference (w/ VLM minus w/o VLM). (b) Corresponding analysis across 24 Tabula Sapiens tissues. Blue and red dots indicate scores with and without VLM, respectively.

**Supplementary Fig. 9.**
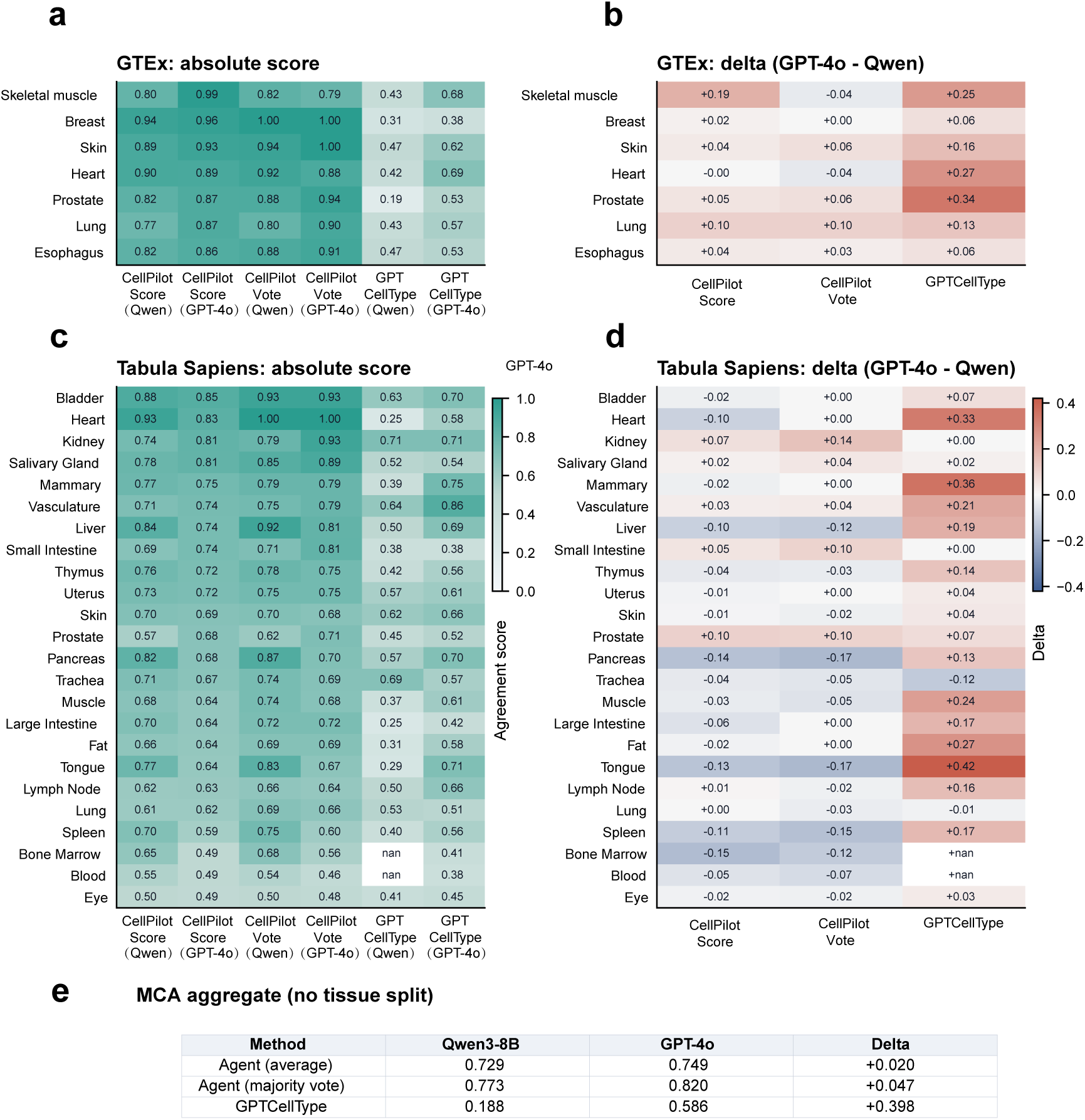
Backbone sensitivity of CellPilot and GPTCelltype across single-cell benchmarks. (a)(c) Absolute agreement scores for GTEx and Tabula Sapiens tissues, respectively, comparing CellPilot with Qwen3-8B and GPT-4o backbones under average score and majority-vote evaluation, and GPTCelltype with the same two backbones. **b, d,** Corresponding GPT-4o minus Qwen3-8B differences for GTEx and Tabula Sapiens. **e,** Aggregate MCA results without tissue-level stratification. CellPilot maintains high agreement with relatively small backbone-dependent changes, whereas GPTCelltype shows larger gains when moving from Qwen3-8B to GPT-4o, particularly in several Tabula Sapiens tissues and in the MCA benchmark. These results indicate that the structured agent framework reduces dependence on model scale while preserving strong annotation performance across datasets.

**Supplementary Fig. 10.**
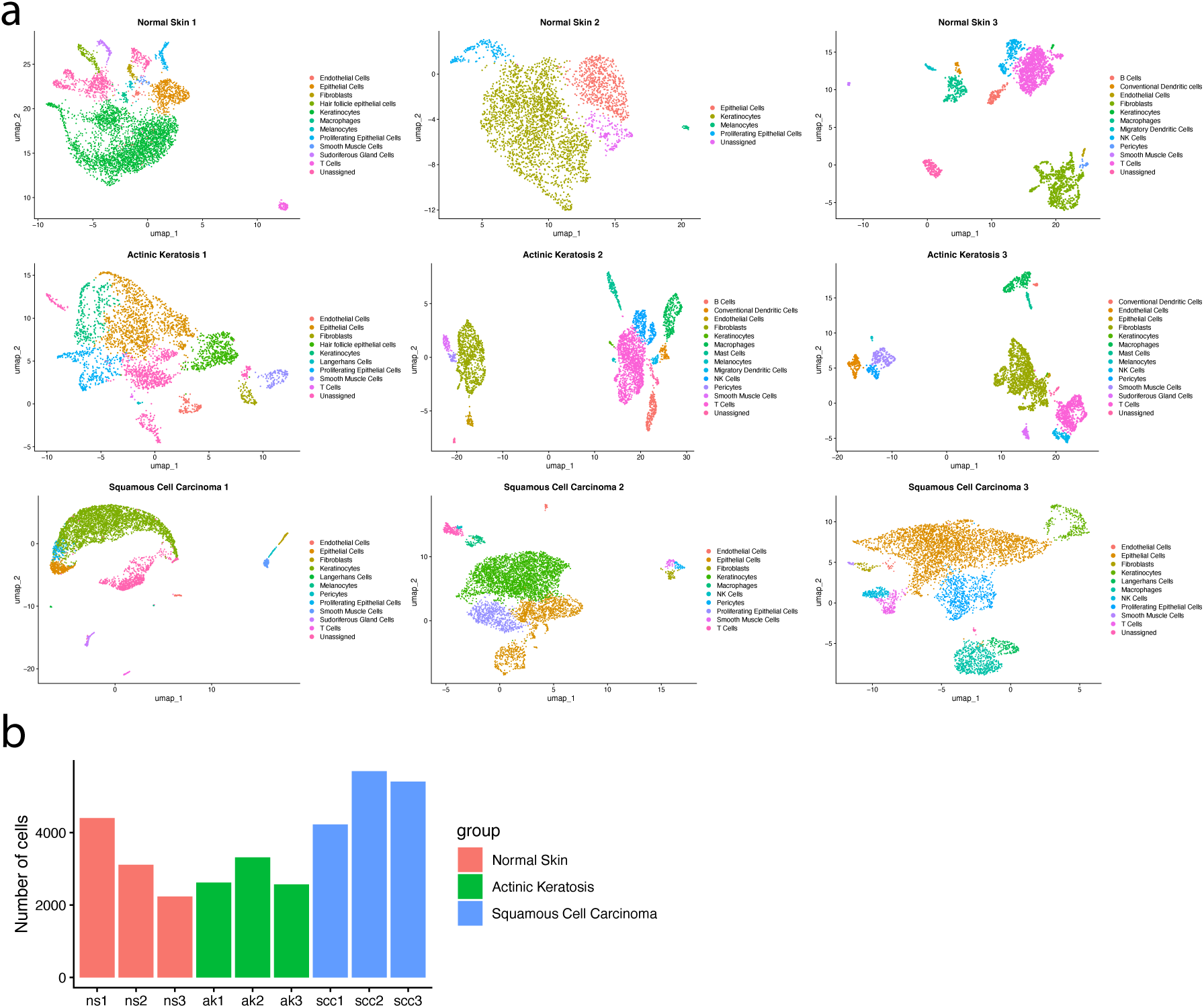
Sample-wise UMAP visualization of agent-generated cell type annotation. (a) Uniform Manifold Approximation and Projection (UMAP) embeddings of single-cell transcriptomes from nine individual skin samples. Each panel represents one sample, with cells colored according to cell type annotations generated by the agent. (b) Bar plot showing the total number of cells profiled in each sample, grouped by condition.

**Supplementary Fig. 11.**
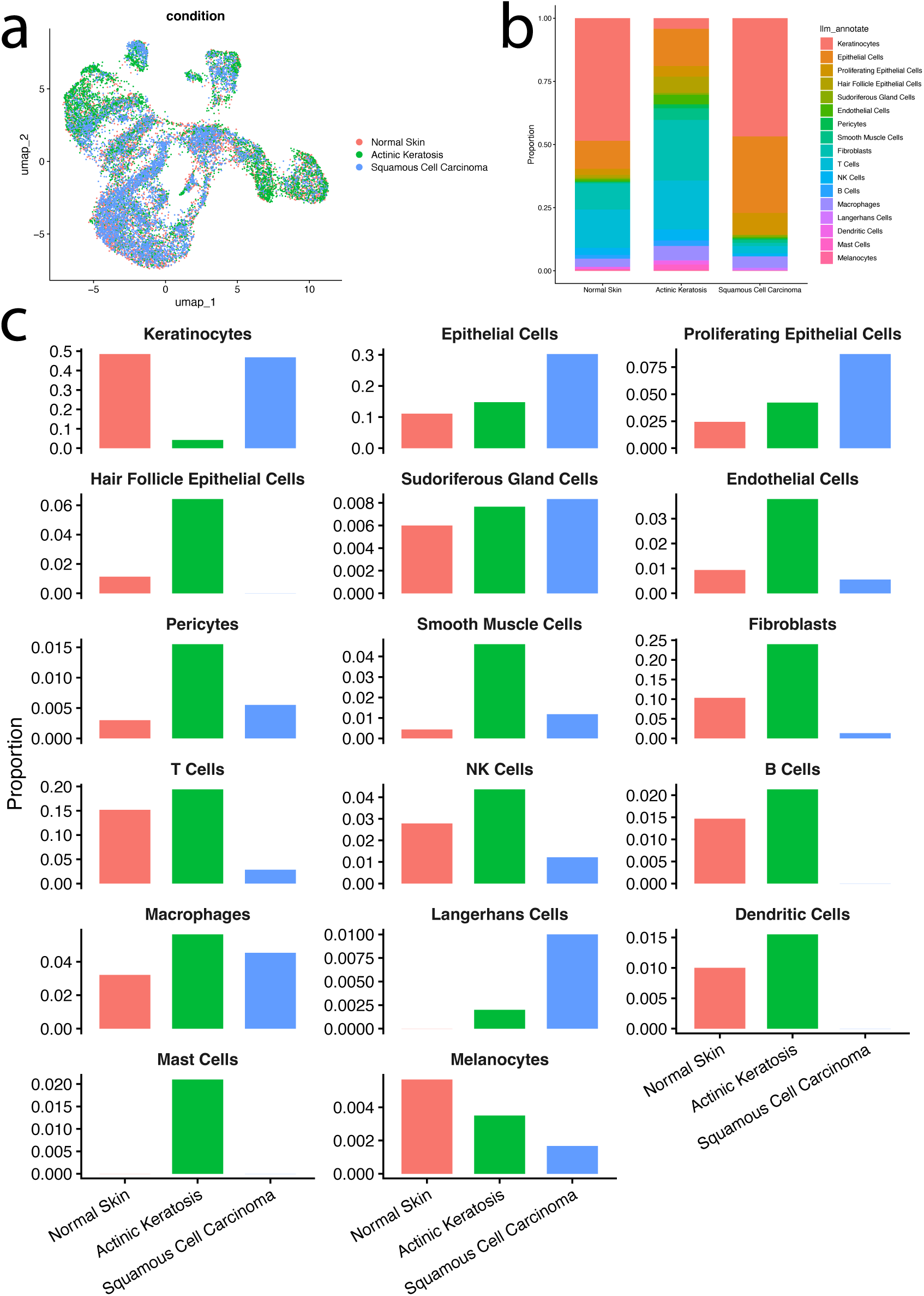
Cell-type proportion changes across normal skin, actinic keratosis and squamous cell carcinoma. (a) Uniform Manifold Approximation and Projection (UMAP) of all cells integrated across nine samples, colored by disease condition (normal skin, actinic keratosis, and squamous cell carcinoma). (b) Stacked bar plot showing the proportion of annotated cell types within each condition. (c) Bar plot showing cell-type specific proportion changes across disease conditions.

